# Identifying summary statistics for approximate Bayesian computation in a phylogenetic island biogeography model

**DOI:** 10.1101/2023.10.13.562305

**Authors:** Shu Xie, Luis Valente, Rampal S. Etienne

## Abstract

Estimation of parameters of evolutionary island biogeography models, such as colonization and diversification rates, is important for a better understanding of island systems. A popular statistical inference framework is likelihood-based estimation of parameters using island species richness and phylogenetic data. Likelihood approaches require that the likelihood can be computed analytically or numerically, but with the increasing complexity of island biogeography models, this is often unfeasible. Simulation-based estimation methods may then be a promising alternative. One such method is approximate Bayesian computation (ABC), which compares summary statistics of the empirical data with the output of model simulations. However, ABC demands the definition of summary statistics that sufficiently describe the data, which is yet to be explored in island biogeography. Here, we propose a set of summary statistics and use it in an ABC framework for the estimation of parameters of an island biogeography model, DAISIE (Dynamic Assembly of Island biota through Speciation, Immigration and Extinction). For this model, likelihood-based inference is possible, which gives us the opportunity to assess the performance of the summary statistics. DAISIE currently only allows maximum likelihood estimation (MLE), so we additionally develop a likelihood-based Bayesian inference framework using Markov Chain Monte Carlo (MCMC) to enable comparison with the ABC results (i.e., making the same assumptions on prior distributions). We simulated phylogenies of island communities subject to colonization, speciation, and extinction using the DAISIE simulation model and compared the estimated parameters using the three inference approaches (MLE, MCMC and ABC). Our results show that the ABC algorithm performs well in estimating colonization and diversification rates, except when the species richness or amount of phylogenetic information from an island are low. We find that compared to island species diversity statistics, summary statistics that make use of phylogenetic and temporal patterns (e.g., the number of species through time) significantly improve ABC inference accuracy, especially in estimating colonization and anagenesis rates, as well as making inference converge considerably faster and perform better under the same number of iterations. Island biogeography is rapidly developing new simulation models that can explain the complexity of island biodiversity, and our study provides a set of informative summary statistics that can be used in island biogeography studies for which likelihood-based inference methods are not an option.

## Introduction

Islands provide great opportunities for evolutionary and biogeographical studies. Their geographical isolation makes it easier to make inferences on their unique ecological and evolutionary processes, because of their well-defined boundaries. It also promotes endemism, providing opportunities to study speciation and diversification. The field of island biogeography is particularly concerned with understanding how island characteristics shape colonization, extinction and speciation rates, and how these affect species richness on islands (MacArthur & Wilson, 1963; Warren et al., 2015). Present-day island biogeography integrates ecological dynamics with evolutionary processes, and considers the disequilibrium of the islands affected by geological ontogeny over time (Whittaker et al., 2008; 2017). In recent years, advances in molecular biology and computational methods have allowed researchers to incorporate phylogenetic information into evolutionary models (Rabosky and Glor 2010; Ronquist, 2011; Valente et al., 2015), allowing a greater understanding of the evolutionary history and relationships in island biotas. Key parameters in island biogeography that can potentially be estimated from island phylogenetic and diversity data include rates of dispersal, colonization, speciation, extinction, island carrying capacity, the strength of ecological interactions, as well as rates of co-evolution and trait evolution (Patiño et al., 2017). However, the application of phylogenetic information to estimate parameters in island biogeography has remained relatively rare (Lim & Marshall, 2017; Valente et al., 2018).

Accurately estimating rates of colonization and diversification (speciation minus extinction) in island biogeography models is essential for testing hypotheses on the macroevolution and biogeography of island species and predicting future patterns. Maximum likelihood and Bayesian estimation are widely used approaches for parameter inference in a phylogenetic context (Ellison 2004; Sanmartín et al. 2008). Maximum likelihood estimation is asymptotically unbiased and efficient, and also allows for straightforward model comparison (Akaike, 1973). However, the estimation approach can be sensitive to the choice of initial values, which could lead to local rather than global likelihood maxima (Rogers and Swofford, 1999). Bayesian inference is an alternative probabilistic approach that requires specifying a prior probability distribution reflecting the prior knowledge of the parameters. The posterior distribution is updated using Bayes’ theorem incorporating the likelihood and the prior. Compared with a single point estimation obtained through maximum likelihood, Bayesian inference can be more informative, providing a distribution of possible parameter values, and allowing for an escape from local maxima. Markov Chain Monte Carlo (MCMC) is a typical method used in the Bayesian inference framework. MCMC has been particularly developed for cases when only the likelihood ratio or the likelihood of an augmented data set is available, but here we use it with the full likelihood function. While efficiently computing the likelihood or likelihood ratio can be done for simple models of island biodiversity dynamics, the increasing complexity of models in island biogeography and the resulting increase in parameters to be estimated makes it unfeasible. Thus, there is a need for likelihood-free parameter estimation methods.

A promising approach to overcome this challenge is approximate Bayesian computation (ABC), a likelihood-free simulation-based Bayesian method. The ABC algorithm estimates model parameters by simulating data, and accepts the parameters which can generate simulations close enough to the observations according to summary statistics and distance measures (Beaumont et al., 2002; Toni & Stumpf, 2009). The ABC approach originated in the context of population genetic analysis with a basic ABC rejection algorithm (Beaumont, 2010). However, this algorithm can be inefficient with a low acceptance rate in complex models, especially when the prior distribution is difficult to specify (Sadegh and Vrugt, 2014). In the past decade, Monte Carlo algorithms, such as Markov chain Monte Carlo (MCMC) or sequential Monte Carlo (SMC), have been incorporated in ABC and have been applied to infer diversification rates in ecological or evolutionary models (Jabot & Chave, 2009; Rabosky, 2009; Bokma, 2010; Slater et al., 2012; Janzen et al., 2015). The ABC-SMC algorithm has become a popular method as it allows for tackling large and complex systems.

The computational efficiency and performance of the ABC algorithm are largely determined by the choice of summary statistics (Janzen et al., 2015; Jiang et al. 2017). In macroevolutionary studies, potentially informative statistics are those describing the species richness and the endemism of the extant species of interest. In addition, summary statistics that quantify the phylogenetic relationships between species and branching times may also be important. In island biogeography, phylogenetic data provide information on whether island species are derived from single or multiple independent colonization events. Furthermore, phylogenies also help to identify the biogeographic regions from which island colonizers originated, and can provide information on the role of dispersal or vicariance in shaping the distributions of island lineages. In a likelihood context, the inference accuracy of island colonization, speciation and extinction rates has been shown to be improved by adding temporal phylogenetic information on the pattern of accumulation of species on island, particularly the colonization times of island lineages, but also cladogenetic branching times (Valente et al. 2018). Hence identifying summary statistics capturing phylogenetic information seems a promising strategy.

Phylogenetic diversity (PD) and mean pairwise distance (MPD) are widely used statistics in phylogenetic analysis (Tsirogiannis & Sandel, 2016). Another promising summary statistic, the normalized Lineage Through Time (nLTT), was proposed to assess the similarity between any two phylogenetic trees (Janzen et al., 2015). It was shown that nLTT performs better than traditionally used statistics (PD, tree size and gamma statistic) in estimating diversification rates in birth-death, time-dependent and diversity-dependent speciation models (Janzen et al., 2015). The good performance of this statistic can be explained by the fact that it has been shown to be a sufficient statistic for the estimation of speciation and extinction rates in a constant-rate or time-dependent birth-death model of diversification (Louca & Pennell, 2020). Furthermore, while PD and MPD statistics are calculated based on binary trees, in which all the species evolved from the same ancestor, (a variant of) the LTT statistic can also be used for analyzing phylogenies of multiple independent clades (Neves et al., 2022). This is relevant in island biogeography models, where the focus is often on the diversity dynamics of the entire island community, that is, the phylogenetic trees composed of the clades descending from multiple independent colonization events.

The ABC approach has been used in island biogeography on population genetic analyses (Patiño et al., 2015), but remains relatively underutilized in macroevolutionary models. Island biogeography is rapidly developing new models that can better explain the complexity of island biodiversity, for example incorporating geodynamics (Borregaard et al 2015, Neves et al., 2022), trait evolution (Xie et al., 2023), or both (Sukumaran et al., 2016). It is thus essential to identify general but efficient summary statistics, particularly for phylogenies of island species and communities, in order to be able to use ABC approaches to estimate parameters in complex island biogeography models (which are not amenable to likelihood methods). In this study, we develop an ABC-SMC framework and test its inference ability on simulations with the island biogeography model, DAISIE (Dynamic Assembly of Island biotas through Speciation, Immigration and Extinction) (Valente et al., 2015) by comparing with the results using likelihood-based methods. DAISIE integrates phylogenetic information into island biogeography. It allows estimation of the rates of colonization, extinction and speciation (CES) from phylogenetic data, and can be used to simulate these processes for given rates. The DAISIE simulation framework was expanded into various directions incorporating island ontogeny and sea-level change (Neves et al., 2022) and trait effects (Xie et al., 2023). However, computing the likelihood is often not possible for these extended models, and thus a likelihood-free ABC algorithm is necessary for parameter estimation. To facilitate comparison with the ABC results (i.e., making the same assumptions on prior distributions), here we also develop a likelihood-based Bayesian inference framework using Markov Chain Monte Carlo (MCMC). We simulated a variety of island biogeography scenarios with the DAISIE simulation model and compared the estimated values using the three inference approaches: MLE, ABC and MCMC. We asked the following questions by comparing the estimation performance: 1) Can the ABC algorithm accurately estimate parameters in phylogenetic island biogeography models (as accurately as MLE or MCMC)? 2) If not, under what conditions are estimations poor? 3) Which summary statistics are most relevant and efficient for ABC island biogeography studies? 4) Are phylogenetic summary statistics more informative than statistics of the tips of the tree (e.g. species richness)?

## Methods

### Simulation model and scenarios

Our focal system is a community of extant species living on an island, for which one would like to estimate rates of colonization, speciation and extinction based on diversity and phylogenetic data. Hence, we used as observations island diversity and phylogenetic data simulated in the DAISIE framework. The species richness on the island is determined by the colonization rate γ, anagenesis rate λ_a_, cladogenesis rate λ_c_ and extinction rate μ. Cladogenesis and colonization rates can be diversity-dependent, declining linearly with an increasing number of species on the island. The simulation output is a set of phylogenies of island species each resulting from an independent colonization event of the island. Some phylogenies may have multiple species (e.g. island radiation), and others may have just a single island species (e.g., non-endemic species that have not had time to speciate or endemic singleton species with no close relatives on the island). To mimic empirical studies in which it is difficult to obtain complete extinction information, we pruned all extinct species from the simulations and used as data entries the reconstructed trees containing all species that are extant on the island at the end of the simulation. We used the standard DAISIE simulation code implemented in the R package DAISIE (Etienne et al., 2023). We do not consider extensions of the original model (such as trait-dependence or area-dependence of rates) because there are currently no likelihood methods for these more complex models. The simulated data produced in this way are hereafter referred to as observed data, as they are meant to mimic empirical data, and to distinguish them from the simulations performed in the ABC analysis.

We used a total of 16 parameter combinations (Table 1) to simulate observed islands (composed of one or more phylogenetic trees, each representing a lineage descending from a colonization event). We ran 10 replicates for each parameter combination, thus obtaining 160 simulated islands in total as observed data that were used for the subsequent inference using different estimation methods. We set the island age as 5 million years in all the simulations. We tested the estimation performance in a diversity-independent (DI) model with no upper bound for the species richness on the island.

**Table 1.** Parameter space used to generate the observed data used in ABC-SMC and MCMC algorithms. In total, 16 parameter combinations were used.

| Parameters | Meaning | Parameter values |
| --- | --- | --- |
| $\lambda^c$ | Cladogenetic speciation rate | 0.4, 0.7 |
| $\mu$ | Extinction rate | 0, 0.3 |
| $\gamma$ | Colonization rate | 0.003, 0.009 |
| $\lambda^a$ | Anagenetic speciation rate | 0.1, 1.0 |

### Maximum likelihood estimation

A likelihood computation method already exists in the DAISIE framework (Valente et al., 2015). The method allows estimating colonization, speciation and extinction rates using maximum likelihood, based on the phylogenetic information of colonization and branching times, as well as the endemicity status of species (endemic or not endemic to the island). The code for parameter estimation is implemented in the R package DAISIE (Etienne et al., 2023).

### Metropolis-Hastings MCMC algorithm

We developed and performed a Metropolis–Hastings Markov chain Monte Carlo (MCMC, Metropolis et al. 1953; Hastings 1970) analysis to obtain our reference estimates on the observed data using the analytical likelihood (Valente et al. 2015). In Bayesian MCMC, a sequence (chain) of parameter values is generated using Monte Carlo sampling, where each step in the chain only depends on the previous step (Markov property). The chain is designed such that the distribution of the generated parameter values converges to the posterior distribution of the parameters. We ran 1,000,000 iterations for the MCMC chain, with a discarded burn-in phase of 100,000 at the beginning, and the chain was thinned by sampling every 400th iteration, with low autocorrelation coefficient values (< 0.15). In order to reduce the influence of the prior distribution, we chose uniform distributions as the prior distributions for each parameter, using two sets of prior distributions with different ranges (Table 2) to evaluate the effect of the choice of the prior.

**Table 2.** The ranges of the two uniform prior distributions used in ABC-SMC and MCMC algorithms.

| Parameters | Prior distribution 1 | Prior distribution 2 |
| --- | --- | --- |
| $\lambda^c$ | [0, 1] | [0, 2] |
| $\mu$ | [0, 0.5] | [0, 2] |
| $\gamma$ | [0, 0.02] | [0, 0.02] |
| $\lambda^a$ | [0, 1.5] | [0, 2] |

### ABC-SMC algorithm

We applied a likelihood-free ABC-SMC algorithm to the observed data to estimate parameters (Toni et al., 2009). ABC-SMC uses a sequential sampling approach that generates a series of intermediate distributions with narrowing-down scales (Fig 1). In our ABC-SMC algorithm, we first sample a group of parameters (collectively called *particle*) from the prior and perturb the parameters using a normal distribution with zero mean and a standard deviation of 0.05, and then we use these to simulate islands with the DAISIE simulation model. The similarity between the observed data and the simulated data is assessed with the distance between vectors of summary statistics. The particles are accepted when the distance for each summary statistic between the two datasets is smaller than a predefined threshold ε. Otherwise, the particle should be resampled from the prior distribution until accepted. When a sufficient number of particles (set beforehand at 500 in our simulations) is accepted, then a new iteration starts, where again particles are sampled, but now from the previous parameter population. Then the above processes of simulating datasets and accepting particles is repeated (Fig 1). The threshold decreases with each new iteration (ε_1_ > ε_2_ >… > ε_n_), where we use the median values of the summary statistic distances as the new thresholds in the following iteration. The algorithm stops when the acceptance rate is lower than 0.002 (one accepted particle in every 500 simulation steps), and we assume that the particles have converged. The population of the samples in the last iteration are regarded as the posterior, which is used to compare with the other approaches.

**Figure 1.**
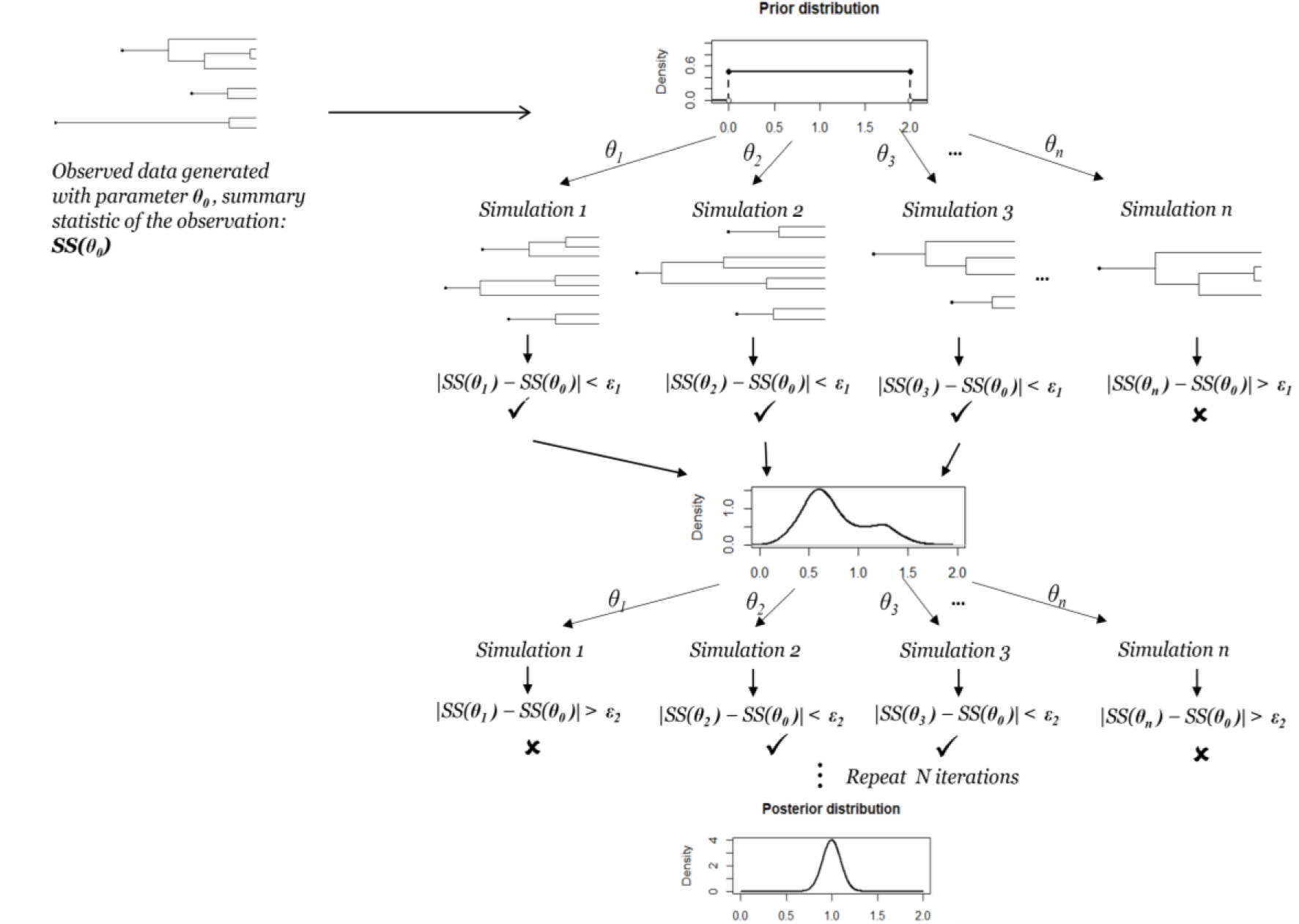
Illustration of the ABC-SMC algorithm. *θ_1_*, *θ_2_*,…, *θ_n_* indicate n particles sampled from the prior distribution, and *SS(θ)* is the summary statistic of the particle (a single simulation). *ε* is the distance threshold for each iteration, which decreases with each iteration.

To be comparable, we used the same observed data and the same prior distributions as in the MCMC algorithm (Table 2). In the ABC simulations, we kept the same island age (5 million years) as in the observed island dataset, but the number of species and the number of colonists present on the island can vary stochastically. We also pruned extinct species from each ABC-simulated island dataset.

We selected eight summary statistics that describe the number of extant species on islands, the phylogenetic relationships between species as well as the timing of accumulation of species (times of colonization and branching times) (Table 3). We evaluated the ABC performance using four groups of summary statistics, a) ABC diversity, with three statistics with island species richness and endemism information but without information on evolutionary history: the total number of extant species on the island (*N*_total_), the number of endemic species (*N*_end_), and the number of non-endemic species (*N*_non-end_); b) ABC NLTT, with three LTT statistics with temporal information: the total number of species on the island through time (NLTT_total_), the number of singleton endemic species through time (NLTT_singleton-end_), the number of non-endemic species through time (NLTT_non-end_); c) ABC phylogenetic, with five statistics consisting of the three temporal statistics in the ABC NLTT group plus two additional phylogenetic statistics: the standard deviation of colonization time (SD-CT), and the standard deviation of clade size (SD-CS); d) ABC all, the combination of all eight statistics. Phylogenetic information allows us to determine whether an endemic island clade is composed of a single endemic species (singleton endemic) or of multiple endemic species (radiation). In the three groups of summary statistics that use phylogenetic information (ABC NLTT, ABC phylogenetic and ABC all), we used the NLTT of singleton endemic species (NLTT_singleton-end_) instead of NLTT of all the endemics, because it is more informative on anagenetic speciation (e.g., if there are fewer endemic singleton clades than endemic clades with multiple species, this suggests that cladogenesis is more frequent than anagenesis). We do not lose information, because the number of endemics in a radiation can be calculated from the total number of species, the number of non-endemics and the number of singleton endemics. We calculated NLTT statistics using the R package *nLTT* on CRAN, which is also available on GitHub (https://github.com/thijsjanzen/nLTT). The code used to perform the MCMC and the ABC-SMC analyses was implemented in the R package *DivABC* on GitHub (https://github.com/xieshu95/DivABC).

**Table 3.** Summary statistics used for the ABC algorithm. We consider statistics 1-3 to be diversity-related statistics (ABC diversity), 4-6 to be temporal/phylogenetic statistics (ABC NLTT), and 7-8 to be phylogenetic statistics (ABC phylogenetic).

|  | Summary statistics | Description | Category |
| --- | --- | --- | --- |
| 1 | $N_{\text{total}}$ | Total number of extant species on the island | Diversity-related |
| 2 | $N_{\text{end}}$ | Number of endemic island species | Diversity-related |
| 3 | $N_{\text{non-end}}$ | Number of non-endemic island species | Diversity-related |
| 4 | $\text{NLTT}_{\text{total}}$ | Number of island lineages through time | Phylogenetic (NLTT) |
| 5 | $\text{NLTT}_{\text{singleton-end}}$ | Number of singleton island endemics through time | Phylogenetic (NLTT) |
| 6 | $\text{NLTT}_{\text{non-end}}$ | Number of non-endemic island species through time | Phylogenetic (NLTT) |
| 7 | SD-CT | Standard deviation of colonization time among island clades | Phylogenetic |
| 8 | SD-CS | Standard deviation of clade size among island clades | Phylogenetic |

## Results

We compared the performance of the parameter estimation using the MLE, ABC and MCMC approaches. We found that MLE performs the best. The ABC algorithm performs well in estimating the colonization, extinction and speciation (CES) rates when the colonization rate is high, and thus the observed data contain more clades and species on the island. For most of the parameter sets we tested in this study, using phylogenetic statistics in the ABC approach results in higher estimation accuracy than using only diversity-related statistics. Furthermore, for all three methods (MLE, MCMC and ABC), we find that the inference error and the standard deviation decrease as the species richness and the number of clades increase.

### Observed data

The island datasets generated with different parameter sets show large differences in the total species diversity and the number of clades present on the island (Table 4). As for the endemicity status, the number of non-endemics differs less than the number of endemic species (Table S1). High cladogenesis (λ_c_ = 0.7) and colonization (γ = 0.009) rates can cause some islands to have more than 500 extant species in total, with the largest recorded clade containing 129 species (Table S1). The smallest island dataset we obtained in this study has just seven species distributed over five clades, which was obtained under conditions of low colonization (γ = 0.003) and speciation rates (λ_c_ = 0.4), and a high extinction rate (μ = 0.3). Although such highly diverse or very species-poor islands may not be common in empirical studies, we nevertheless wanted to evaluate the inference power in those extreme cases. We calculated the summary statistics of all the observations used in this study, and evaluated the correlations between the statistics with a heatmap (Fig 2). There are strong one-to-one correlations between the diversity-related statistics (N_total_, N_non-end_) and the temporal NLTT statistics (NLTT_total_, NLTT_non-end_), which indicates a high overlap of the information provided by these statistics. N_end_ is independent from NLTT_singleton-end_, but is positively correlated with N_total_. However, SD-CT, which is a statistic to measure the colonization time variance, is independent of other statistics, as is evident from a low correlation coefficient (Fig 2).

**Table 4.** Statistics of the observed datasets. We show the mean (standard deviation) of the 10 replicates for each parameter set.

| Scenario | $\lambda^c$ | $\mu$ | $\gamma$ | $\lambda^a$ | Number of clades | Total species richness | Largest clade |
| --- | --- | --- | --- | --- | --- | --- | --- |
| 1 | 0.4 | 0 | 0.003 | 0.1 | 14 (4) | 44 (15) | 13 (5) |
| 2 | 0.7 | 0 | 0.003 | 0.1 | 14 (3) | 134 (58) | 52 (28) |
| 3 | 0.4 | 0.3 | 0.003 | 0.1 | 9 (3) | 19 (10) | 7 (3) |
| 4 | 0.7 | 0.3 | 0.003 | 0.1 | 11 (5) | 47 (21) | 17 (7) |
| 5 | 0.4 | 0 | 0.009 | 0.1 | 44 (6) | 138 (30) | 20 (9) |
| 6 | 0.7 | 0 | 0.009 | 0.1 | 41 (8) | 352 (124) | 56 (22) |
| 7 | 0.4 | 0.3 | 0.009 | 0.1 | 30 (5) | 64 (16) | 11 (4) |
| 8 | 0.7 | 0.3 | 0.009 | 0.1 | 32 (6) | 159 (41) | 35 (17) |
| 9 | 0.4 | 0 | 0.003 | 1 | 17 (4) | 52 (18) | 14 (9) |
| 10 | 0.7 | 0 | 0.003 | 1 | 16 (2) | 127 (40) | 42 (19) |
| 11 | 0.4 | 0.3 | 0.003 | 1 | 8 (3) | 15 (6) | 5 (2) |
| 12 | 0.7 | 0.3 | 0.003 | 1 | 13 (3) | 55 (22) | 16 (7) |
| 13 | 0.4 | 0 | 0.009 | 1 | 42 (6) | 155 (39) | 22 (13) |
| 14 | 0.7 | 0 | 0.009 | 1 | 44 (6) | 389 (124) | 71 (28) |
| 15 | 0.4 | 0.3 | 0.009 | 1 | 30 (4) | 60 (15) | 9 (4) |
| 16 | 0.7 | 0.3 | 0.009 | 1 | 32 (5) | 152 (47) | 27 (10) |

**Figure 2.**
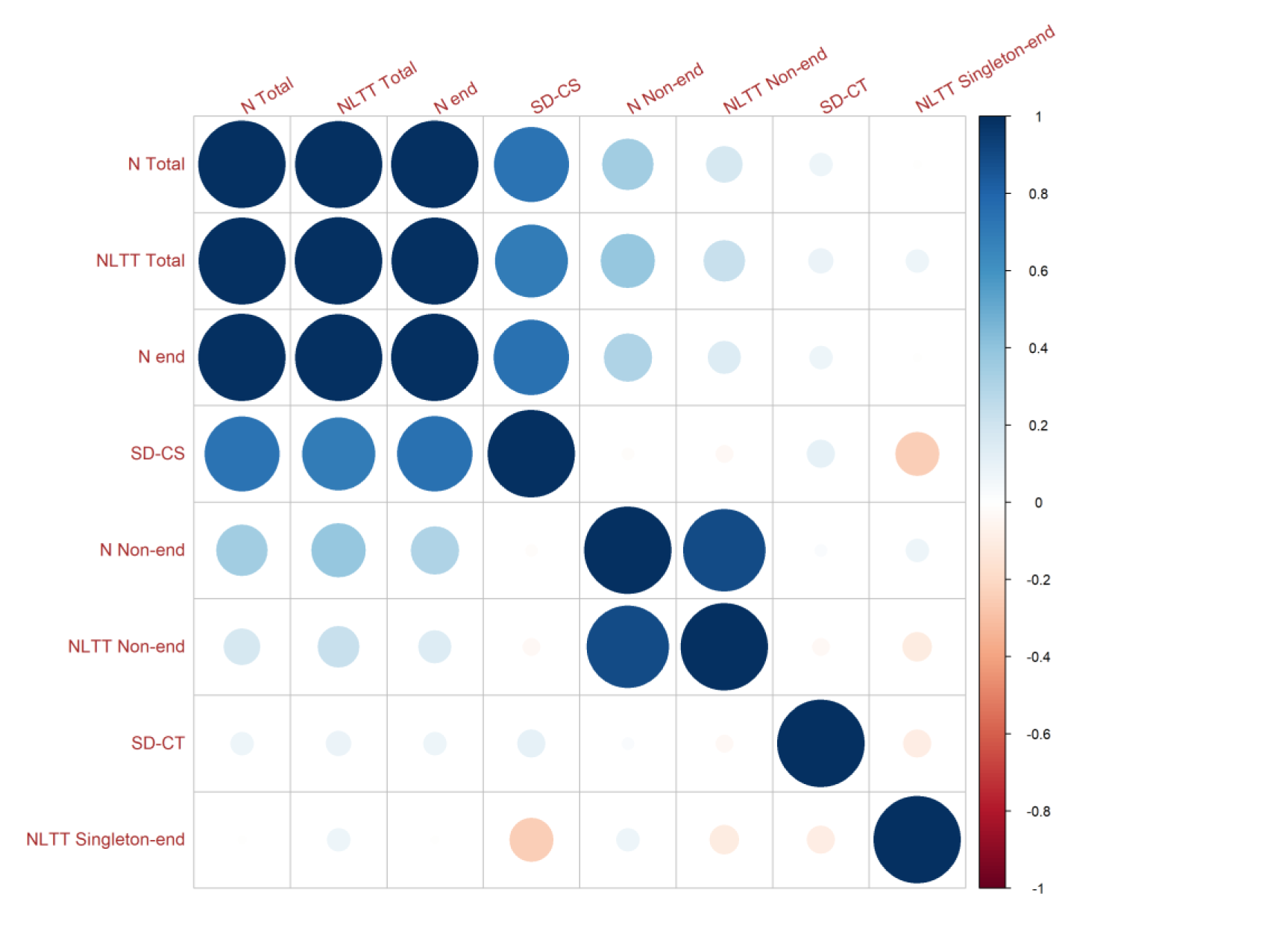
Correlation heatmap between summary statistics of all the observations that were used in the ABC algorithm. Red indicates a negative correlation between two statistics, and blue a positive correlation. The size and color of the circle show the strength of the correlation.

### Inference comparison between ABC, MCMC and MLE

To facilitate comparison between the methods across various parameter combinations used as generating values, we calculated the relative difference between the generating (“true”) rates and the estimated rates, and compared the differences (Δ-rate) between methods. As for the choice of the prior distribution, we find that the ABC estimation performs better under the narrower prior (Fig S1 and S2), with relatively less bias and variance than under the broader prior (Fig 4 and 5), but with the same bias patterns (overestimation or underestimation). Because speciation and extinction rates can often be high on islands (Warren et al., 2015), and because researchers are unlikely to have sufficient information to specify narrow primers, here we discuss the parameter estimations under the broader prior distribution, and we show the estimation results under the narrow prior distribution in supplementary Figure S1 and S2.

Overall, the MLE approach showed the highest accuracy in estimating parameters in the DAISIE framework (Figs 3 and 5), with median Δ values closest to zero for most of the parameter sets. The MCMC and the ABC estimations (except when using diversity-related statistics) have similar performance with minor difference, but both show relatively larger bias in estimating cladogenesis and extinction rates when the generating rate of colonization is low (Fig 5). In addition, because the MCMC and the ABC are Bayesian algorithms producing a distribution as posterior, the standard errors are relatively larger than for the MLE.

**Figure 3.**
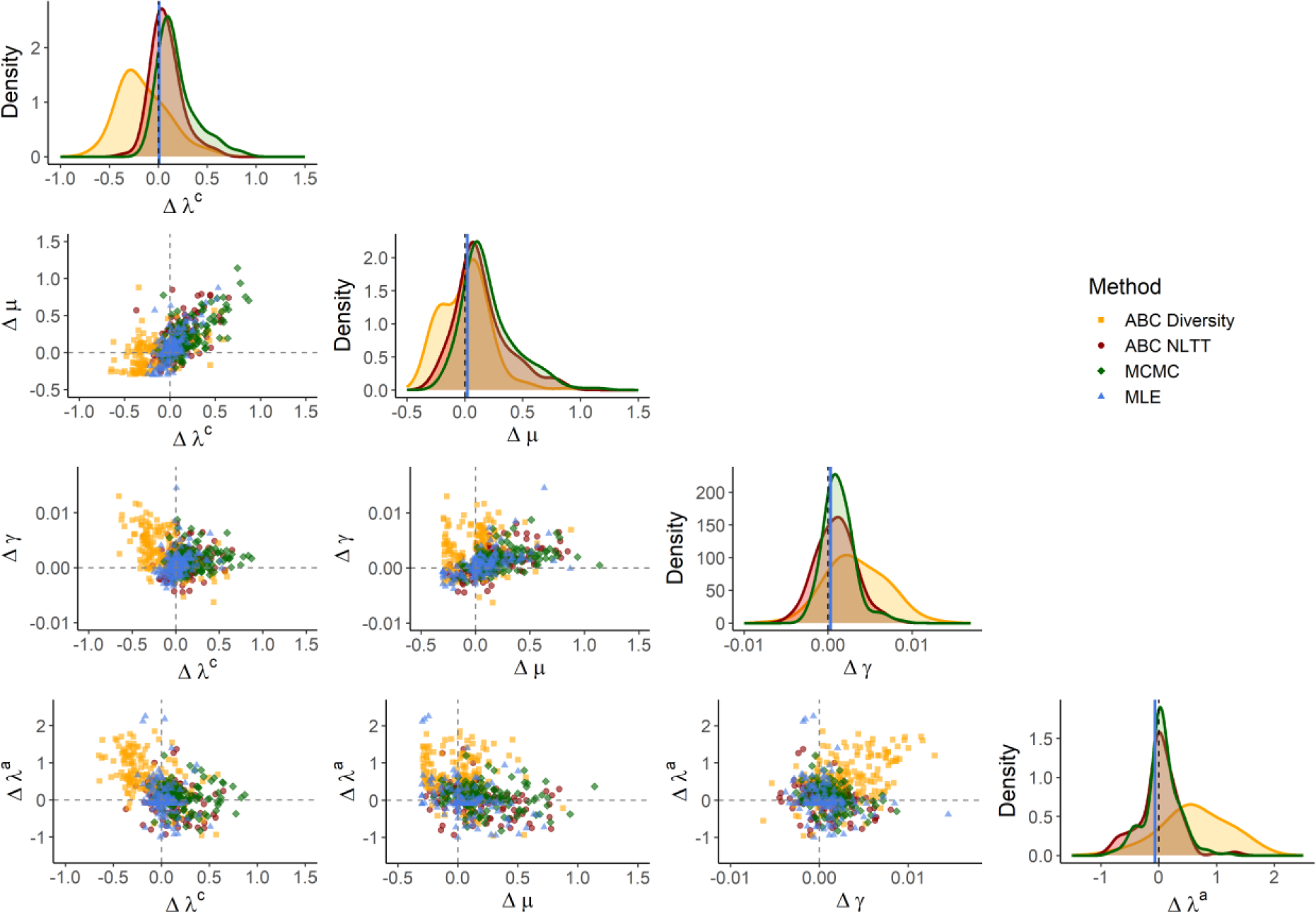
Inference of **cladogenesis, extinction, colonization and anagenesis rate** parameters across all 160 parameter sets. The ABC results are from the analyses using the **broader prior** (See Table 2). Plots show the Δ-rate, that is, the difference between the estimated value and the true value used for generating the observed simulations. The diagonal shows density distributions of the estimations using MCMC (green), ABC with diversity-related (orange) or NLTT (red) statistics. The dashed line is the baseline at zero, and the blue solid line is the median of all MLE estimations across all the parameter sets. The scatter plots in the lower triangle show the Δ-rates for different parameter combinations: the point estimation of MLE (blue triangles), the median value of the posterior distribution using MCMC (green diamonds) and ABC (orange squares and red circles). The gray dashed lines are the baseline, which intersect at zero. ABC Diversity - three diversity-related statistics; ABC NLTT - three NLTT statistics; MLE - maximum likelihood estimation; MCMC - Markov Chain Monte Carlo; γ - colonization rate; μ - extinction rate; λ^c^ – cladogenesis rate; λ^a^ – anagenesis rate.

### Summary statistics

When comparing the posterior distribution of the last ABC iteration we find that, overall, the estimation using phylogenetic statistics (ABC-phylogenetic and ABC-NLTT) appears to be more accurate and stable than that using diversity-related statistics (ABC-diversity), especially when estimating colonization and anagenesis rates (Figs 3-5). We do note that the ABC algorithm with phylogenetic statistics tends to overestimate the cladogenesis and extinction rates when the colonization rate is low (γ = 0.003) (Fig 5). The bias in the extinction rate is larger than that of the cladogenesis rate, therefore resulting in underestimated net diversification rates (Fig 4). The estimations of anagenesis and colonization are accurate for most of the parameter scenarios (Fig 5).

**Figure 4.**
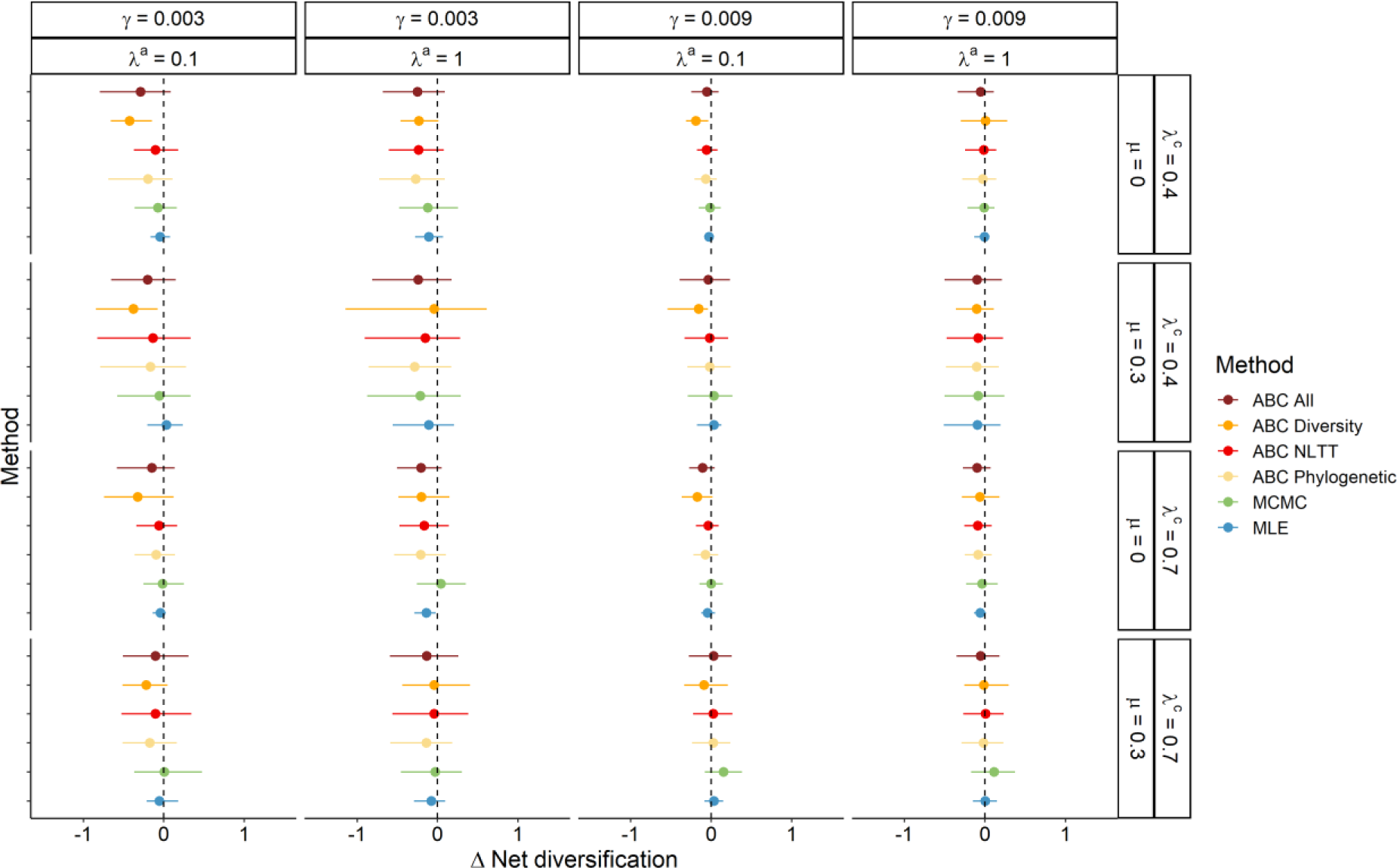
Results of inference of the **net diversification** rate (rate of cladogenesis minus rate of extinction) using MLE, MCMC and ABC. Lines show 95% confidence intervals, dots the median value. The ABC results are from the analyses using the **broader prior** (See Table 2). Plots show the Δ-net diversification rate, that is, the difference between the estimated value and the true value used for generating the observed simulations. The true values are shown at the top of each column and to the right of each row. The distribution for each method combines the 10 replicates for each parameter set. For MLE, these replicates result in 10 data points, whereas for MCMC and ABC these replicates result in 10 posterior distributions combined. In the case of ABC-SMC, the posterior distribution for each replicate is based on 500 particles from the last iteration step of the algorithm. ABC All - all statistics; ABC Diversity - three diversity-related statistics; ABC NLTT - three NLTT statistics; ABC Phylogenetic - five phylogenetic statistics; MLE - Maximum likelihood estimation; MCMC - Markov Chain Monte Carlo; γ - colonization rate; μ - extinction rate; λ^c^ – cladogenesis rate; λ^a^ – anagenesis rate.

**Figure 5.**
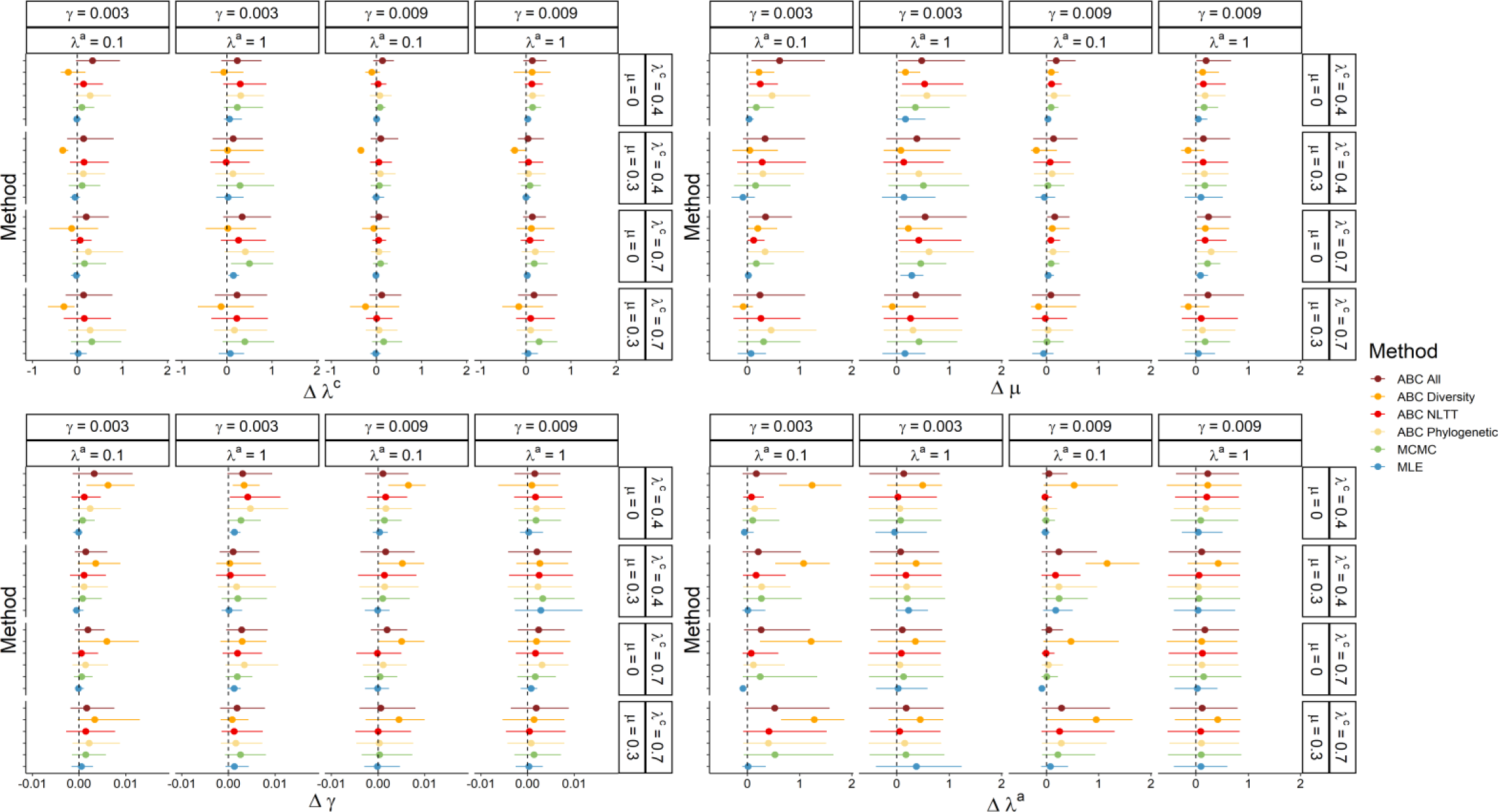
Inference of the **cladogenesis, extinction, colonization and anagenesis rates** using MLE, MCMC and ABC. Lines show 95% confidence intervals, dots the median value. The ABC results are from the analyses using the **broader prior** (See Table 2). Plots show the Δ-rate, that is, the difference between the estimated value and the true value used for generating the observed simulations. The true values are shown at the top of each column and to the right of each row. The distribution for each method combines the 10 replicates for each parameter set. For MLE, these replicates result in 10 data points, whereas for MCMC and ABC these replicates result in 10 posterior distributions combined. In the case of ABC-SMC, the posterior distribution for each replicate is based on 500 particles from the last iteration step of the algorithm. ABC-All – all eight statistics; ABC Diversity – three diversity-related statistics; ABC-NLTT – three NLTT statistics; ABC-Phylogenetic – five phylogenetic statistics; MLE – maximum likelihood estimation; MCMC – Markov chain Monte Carlo; γ – colonization rate; μ – extinction rate; λ^c^ – cladogenesis rate; λ^a^ – anagenesis rate.

The ABC approach using only diversity-related statistics produces large inference bias. Cladogenesis and extinction rates tend to be underestimated, especially when the generating rate of extinction is non-zero (μ = 0.3) (Figs 3 and 5). In addition, colonization and anagenesis rates are largely overestimated for most of the parameter sets (Fig 3), and are more biased when the generating rate of anagenesis is low (λ_a_ = 0.1) (Fig 5). Unlike phylogenetically-informed statistics, diversity-related statistics cannot determine how extant species are divided among clades, or whether an endemic species is in a singleton or in a radiation clade. Therefore, anagenesis rates are difficult to estimate, and tend to be uniformly distributed across the posterior distribution, leading to larger bias. Under the same total number of species and endemic species as in the observed data, an overestimated anagenesis rate may cause more singleton endemic clades and fewer radiations in the ABC simulation (Fig 6), which in turn tends to result in an overestimated colonization rate (Fig 5) and an underestimated net diversification rate. (Fig 4).

**Figure 6.**
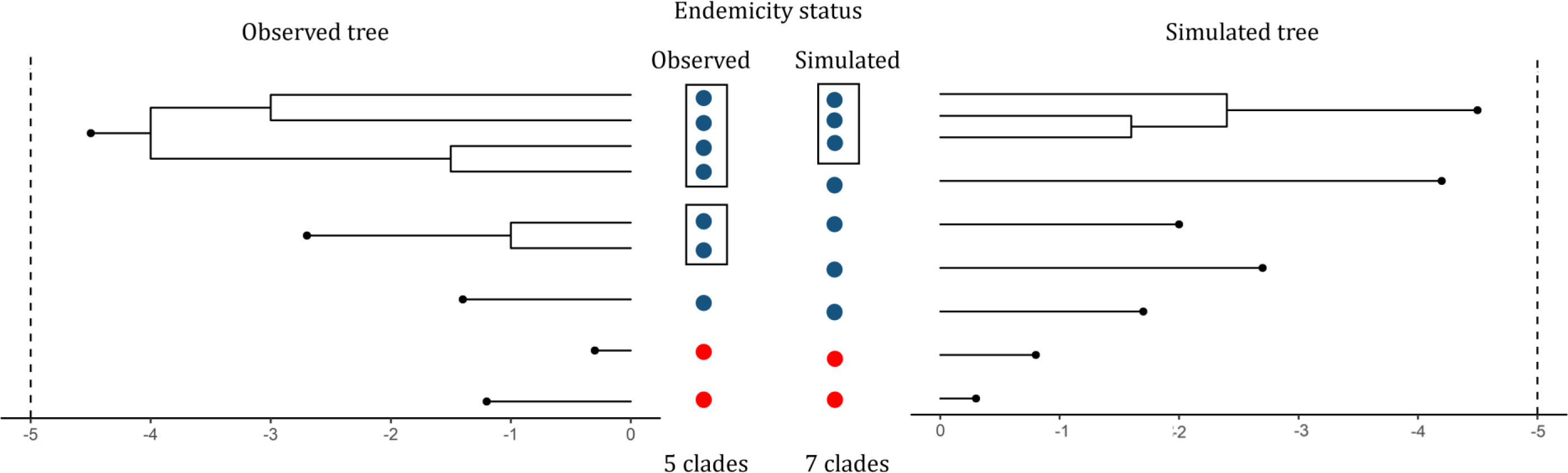
Illustration of how the ABC method with only diversity-related statistics may cause bias in estimating parameters, highlighting the importance of including phylogeny-related statistics. The phylogenetic tree on the left side is an example of an observed dataset, and the tree on the right side is an example of a simulated tree. The two columns in the middle show the endemicity status of the extant species for both observed and simulated data, with non-endemic in red and endemic species in blue. Species inside rectangles are endemic species from the same clade (radiation). The simulated tree has the same total number of species, endemic species and non-endemic species as the observed tree, but a different distribution of species between clades and the number of clades, with many more endemic singletons than the observed data.

If we zoom in on each single replicate (the estimation of each observed dataset), the estimation using diversity-related statistics (ABC-diversity) is more likely to converge to an extremely deviating value (Figs S3 and S4). Furthermore, comparing the estimations under the same iterations, we observe that using phylogenetic statistics always converges faster, which means higher efficiency in estimation (Figs 7 and S5-S8). In addition, estimations with NLTT statistics (ABC-NLTT) contain the least bias and variance comparing the distributions under the same iteration. Adding SD-CS and SD-CT to the three NLTT statistics (ABC-phylogenetic) or combining all the summary statistics (ABC-all) does not provide significant improvement over using only the ABC-NLTT statistics.

**Figure 7.**
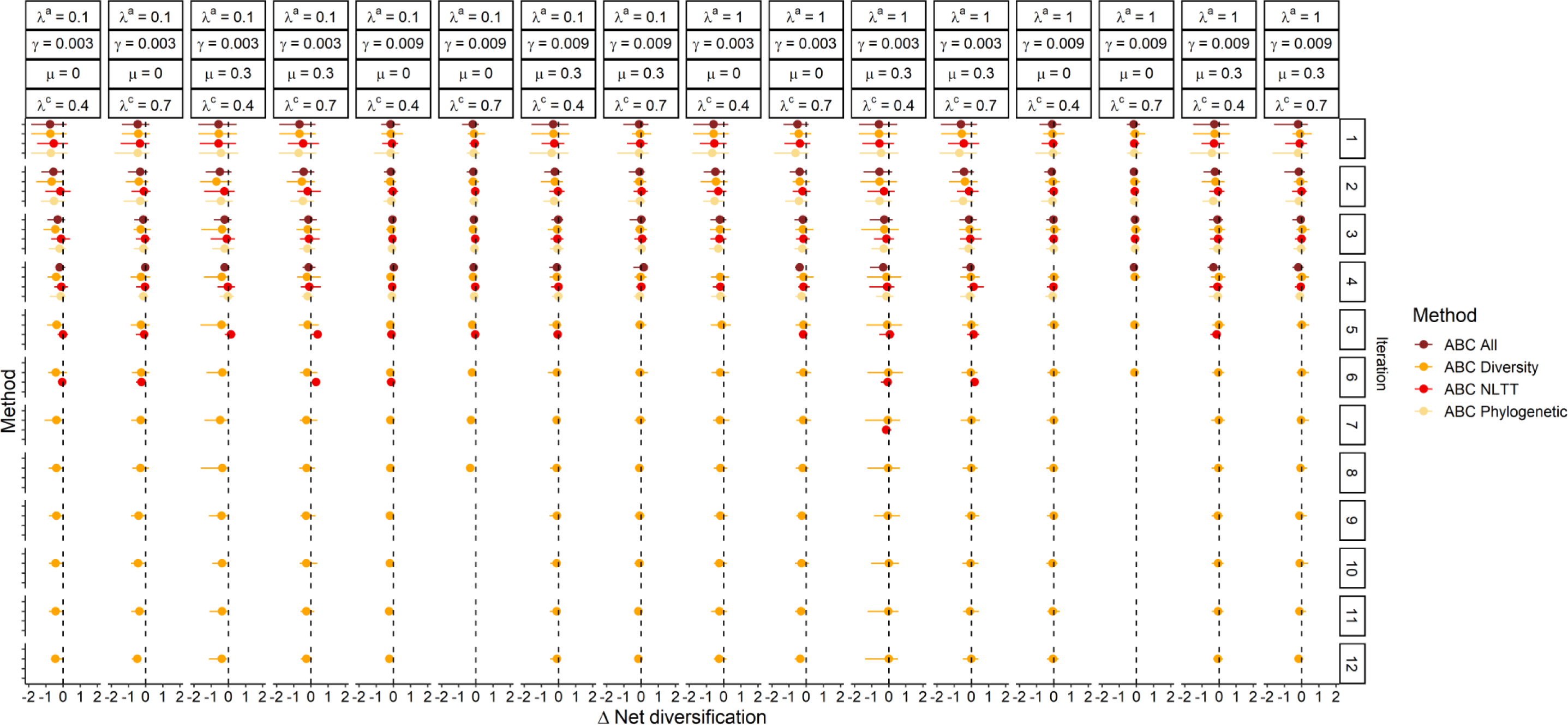
Results of inference of the **net diversification rate** (rate of cladogenesis minus rate of extinction) through **iterations** using ABC with different groups of summary statistics. The ABC results are from the analyses using the broader prior scale (Table 2). Plots show the Δ-net diversification, that is, the difference between the estimated values and the true values used for generating the observed data. The true values are shown at the top of each column. The right of each row shows the number of the ABC iteration. The distribution for each method combines the replicates for each parameter set at a specific iteration. ABC All - all statistics; ABC Diversity - three diversity-related statistics; ABC NLTT - three NLTT statistics; ABC Phylogenetic - five phylogenetic statistics; γ - colonization rate; μ - extinction rate; λ^c^ – cladogenesis rate; λ^a^ – anagenesis rate.

### Correlation between observed dataset properties and estimation

As we found that the generating value of colonization rate plays an important role in estimating parameters, and that colonization rate can affect both the species richness and the number of island clades, we investigated how the parameter estimations depend on properties of the observed data. We used three property metrics to describe the observed datasets: species richness (total number of species on the island), the number of clades, and size of the largest clade. We fitted the relationships between the metrics (median and standard deviation) and the parameter estimations (median) for each observation, and find that all the estimation methods have the same trend: datasets with fewer species and smaller size of the largest clade are likely to cause larger inference and standard errors (Figs 8, and S9, S10 and S11). However, the relationship between the number of clades and the parameter estimations shows two clusters according to the generating colonization rate (Figs 9 and S12). The datasets with higher generating colonization rate contain more clades and have significantly lower bias, especially in estimating cladogenesis and extinction rates.

**Figure 8.**
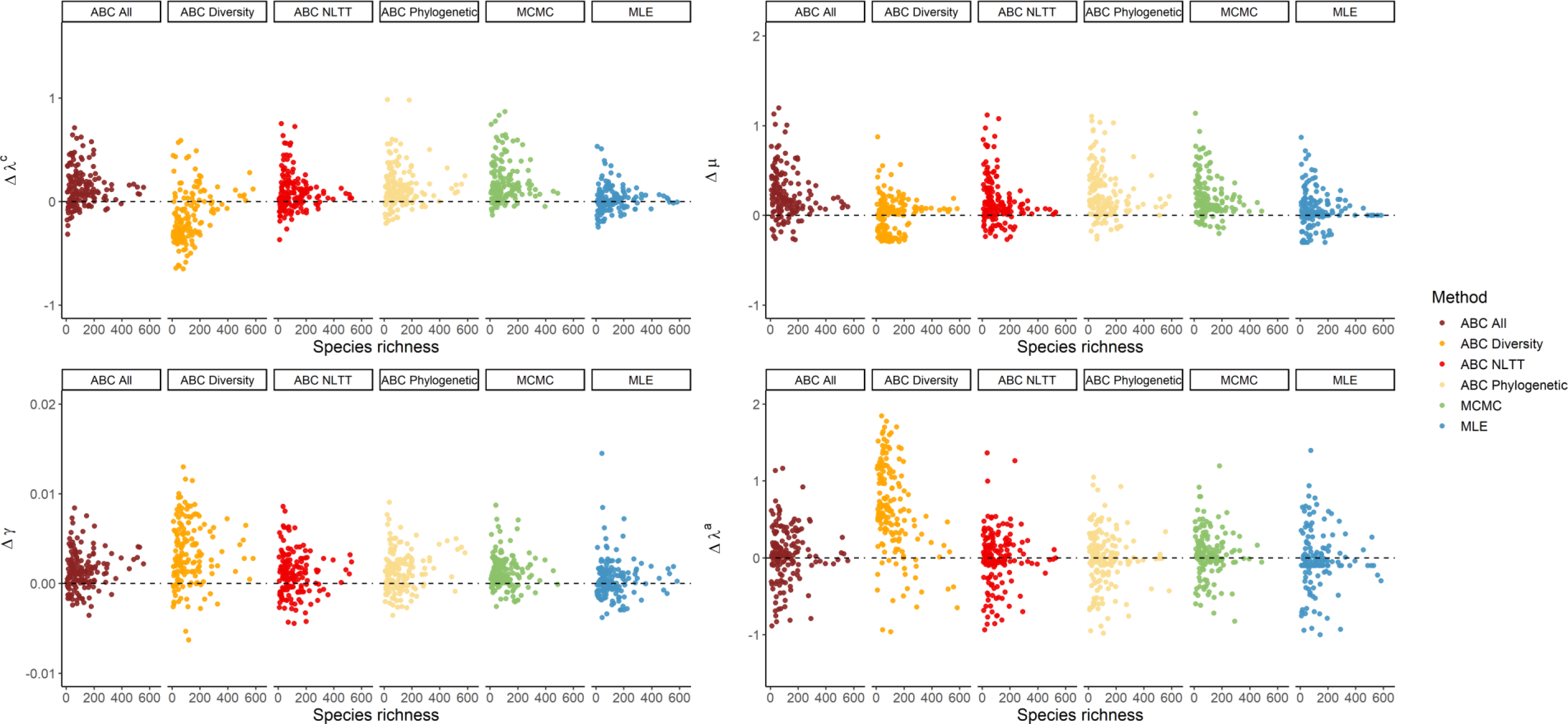
The relationship between the **species richness** of the whole island community and the relative difference in estimating **CES rates**. Different colors indicate different estimation methods. The points show the point estimation of MLE or the **median** value of the posterior distribution using MCMC and ABC.

**Figure 9.**
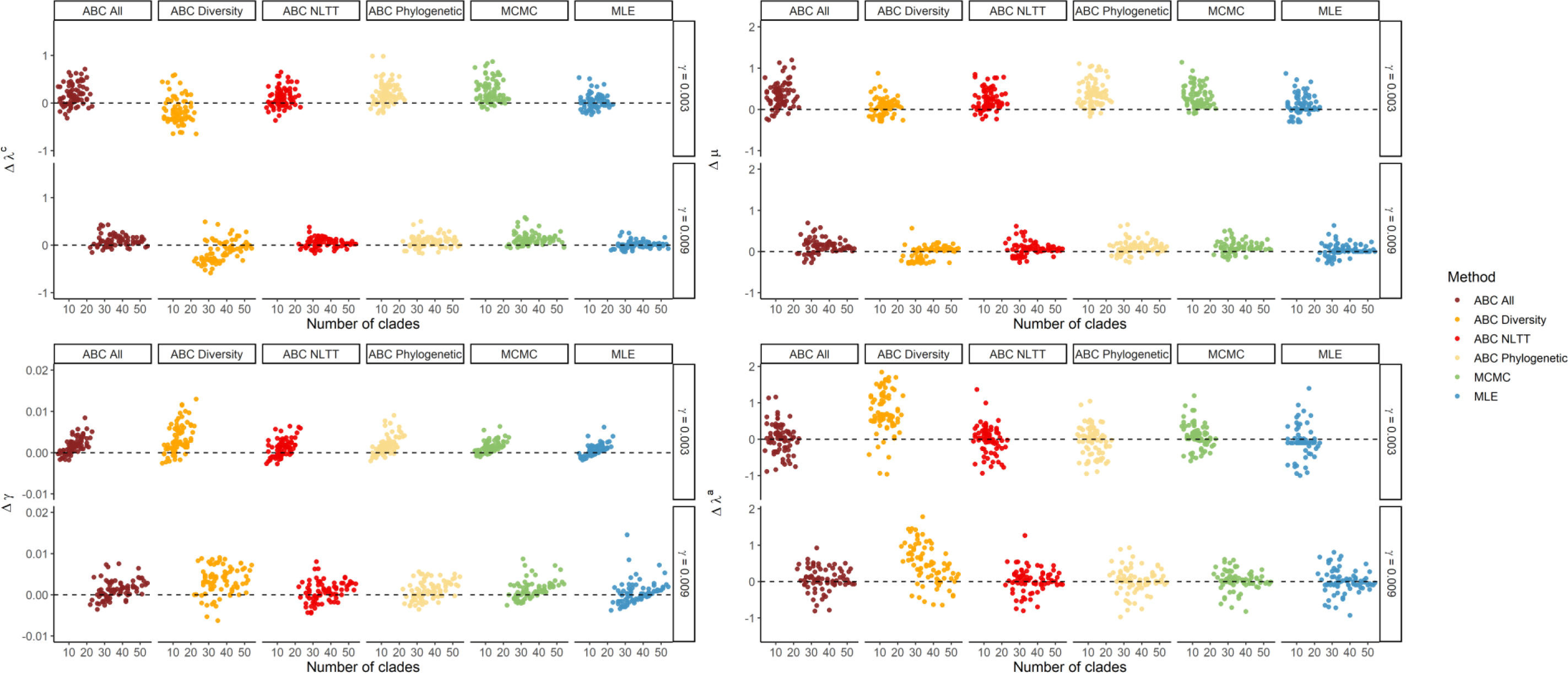
The relationship between **the number of clades** present on the island and the relative difference in estimating **CES rates**. Different colors indicate different estimation methods. The points show the point estimation of MLE or the **median** value of the posterior distribution using MCMC and ABC. The points are clustered based on the generating rates of colonization.

## Discussion

We developed and tested the use of ABC inference in estimating colonization, speciation and extinction rates in an island biogeography model. We compared the estimation performance of the likelihood-free ABC approach with the MCMC and MLE approaches, and evaluated the ABC performance with different sets of summary statistics. Our results indicate that, overall, the performance of the ABC method with phylogenetic statistics is comparable to the likelihood-based methods (especially for MCMC), as the median values of the distributions show minor differences. However, the accuracy of the ABC estimation is sensitive to the generating colonization rate, which is associated with the properties of the observed data (e.g., species richness, the number of clades in the whole island community) (Figs. 8-9 and S9-S12). Furthermore, adding phylogenetic information significantly improves the estimation accuracy and efficiency in the ABC algorithm.

In ABC algorithms, the selection of the prior distribution and the summary statistics significantly affect the power of the estimation (Aeschbacher et al., 2012; Burr & Skurikhin, 2013). We found that a broader prior distribution exacerbates the bias when the observation provides limited information with poor species richness, which was also found in the MCMC results. In this case, higher colonization and cladogenesis than expected (true) rates are more likely to be sampled from broad prior distributions, and can be balanced by high extinction rates to reach the target number of species on the island. However, in the island biogeography context, the extinction or speciation rate could be high in some clades, and from the phylogenetic data one cannot easily extract a suitable prior distribution. Therefore, a broader distribution is necessary to cover the potential true values. In this study, we used uniform distributions as priors. A popular alternative is using the exponential prior distribution for estimating diversification rates (Scarpino et al., 2014). The advantage of using the exponential distribution is to avoid setting boundaries, and always obtain positive values. However, the choice of prior distribution is more important in shaping the posterior when the likelihood function is not highly informative. In this case, using an exponential prior distribution tends to shape the posterior to an asymmetric distribution with low values, which may not be problematic for estimating colonization rates, but could have a significant impact on extinction rates in island biogeography models.

The robustness of the inference on extinction rates is a topic of debate, because of the difficulty in high-precision extinction estimation for single lineages and the lack of information on extinctions (Burin et al., 2019; Louca and Pennell, 2021). However, in the DAISIE framework the likelihood-based approaches can more accurately estimate extinction rates likely because DAISIE works on multiple clades instead of a single lineage. Independent clades can be regarded as multiple replicates, which can be informative for providing almost unbiased estimations (Valente et al., 2015; 2017; 2018). In our study, we also found that the MLE approach can accurately estimate extinction rates for most of the parameter sets (Fig 5). However, because we use pruned phylogenetic trees (both observed and simulations in the ABC algorithm) which exclude extinction events, the ABC algorithm is apparently more sensitive than MLE or MCMC to how much information can be gained from the extant phylogenies (Fig 5), leading to larger inference error in smaller trees. Therefore, the choice of efficient summary statistics, which contain more information on evolutionary dynamics is crucial.

The NLTT statistic is regarded as a powerful statistic that contains evolutionary information as well as the species richness existing at the tips (Janzen et al., 2015; Saulnier et al., 2016; Richter & Wit, 2021). In our study, we find that NLTT statistics are more efficient than diversity-related statistics, with fewer iterations required to reach convergence and less bias (Figs 3-6). In island biogeography, NLTT statistics include the information of colonization time and branching time of each extant clade on the island. When species richness and endemic status are the same, different branching times can lead to variations in NLTT accumulation over time, and the additional phylogenetic information can improve parameter estimation by controlling the temporal variance (Fig 6). This is similar to the conclusions in studies that have shown higher inference accuracy incorporating phylogenetic information (Valente et al., 2018; Antonelli et al., 2018). According to our results, the most efficient combination of the summary statistics in island biogeography modelling is that of NLTT statistics of total species, singleton island endemism and non-endemic species. Including additional statistics does not always result in higher inference accuracy, because they are highly correlated and hence relatively redundant (Fig 2).

The ABC method is considered a good alternative to maximum likelihood estimation, especially when the model is computationally complex. There are a few ABC related R packages implementing ABC algorithms (Csilléry et al., 2012; Jabot et al., 2013), and ABC has been used in a large number of ecological and evolutionary studies (Beaumont, 2010). However, ABC algorithms are rarely used in estimating parameters in macroevolutionary studies in island biogeography. Furthermore, it is rare to find studies comparing the estimation performance between maximum likelihood and ABC when it is possible to analytically calculate the likelihood. This study has identified useful summary statistics to be used for inference in island biogeography models and their extensions. However, more complex summary statistics may be needed for these models. For example, in trait-dependent island biogeography models, trait-related summary statistics will still be required.

