## Supplementary Figures for "Identifying summary statistics for approximate Bayesian computation in a phylogenetic island biogeography model"

Supplementary Materials:

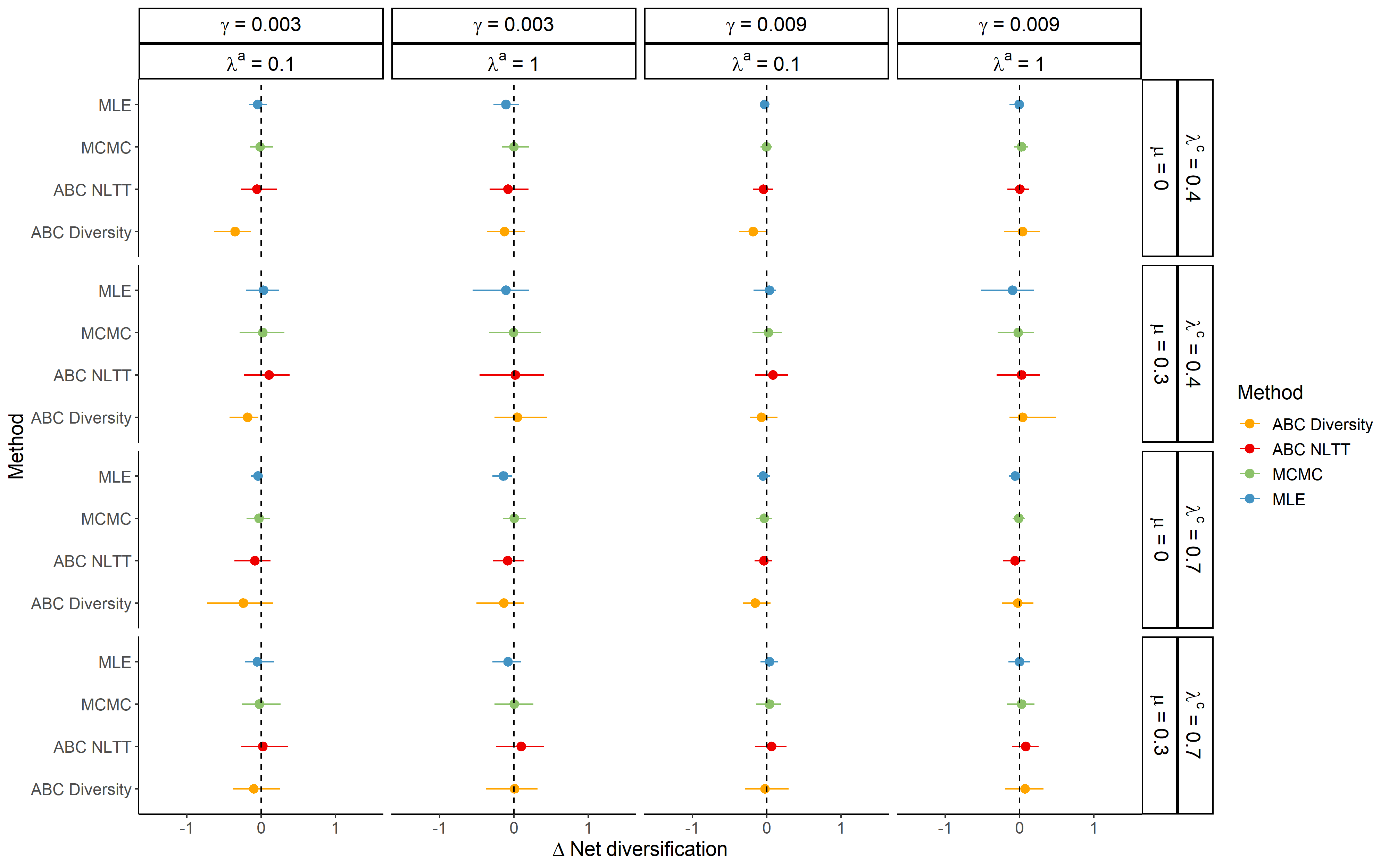

Figure S1. Results of inference of the **net diversification** rate (rate of cladogenesis minus rate of extinction) using MLE, MCMC and ABC. Lines show 95% confidence intervals, dots the median value. The ABC results are from the analyses using the **narrower prior scale** (See Table 2). Plots show the Δ-net diversification rate, that is, the difference between the estimated value and the true value used for generating the observed simulations. The true values are shown at the top of each column and to the right of each row. The distribution for each method combines the 10 replicates for each parameter set. For MLE, these replicates result in 10 data points, whereas for MCMC and ABC these replicates result in 10 posterior distributions combined. In the case of ABC-SMC, the posterior distribution for each replicate is based on 500 particles from the last iteration step of the algorithm. ABC All - all statistics; ABC Diversity - three diversity-related statistics; ABC NLTT - three NLTT statistics; ABC Phylogenetic - five phylogenetic statistics; MLE - Maximum likelihood estimation; MCMC - Markov Chain Monte Carlo; γ - colonization rate; μ - extinction rate; λ^c^ – cladogenesis rate; λ^a^ – anagenesis rate.

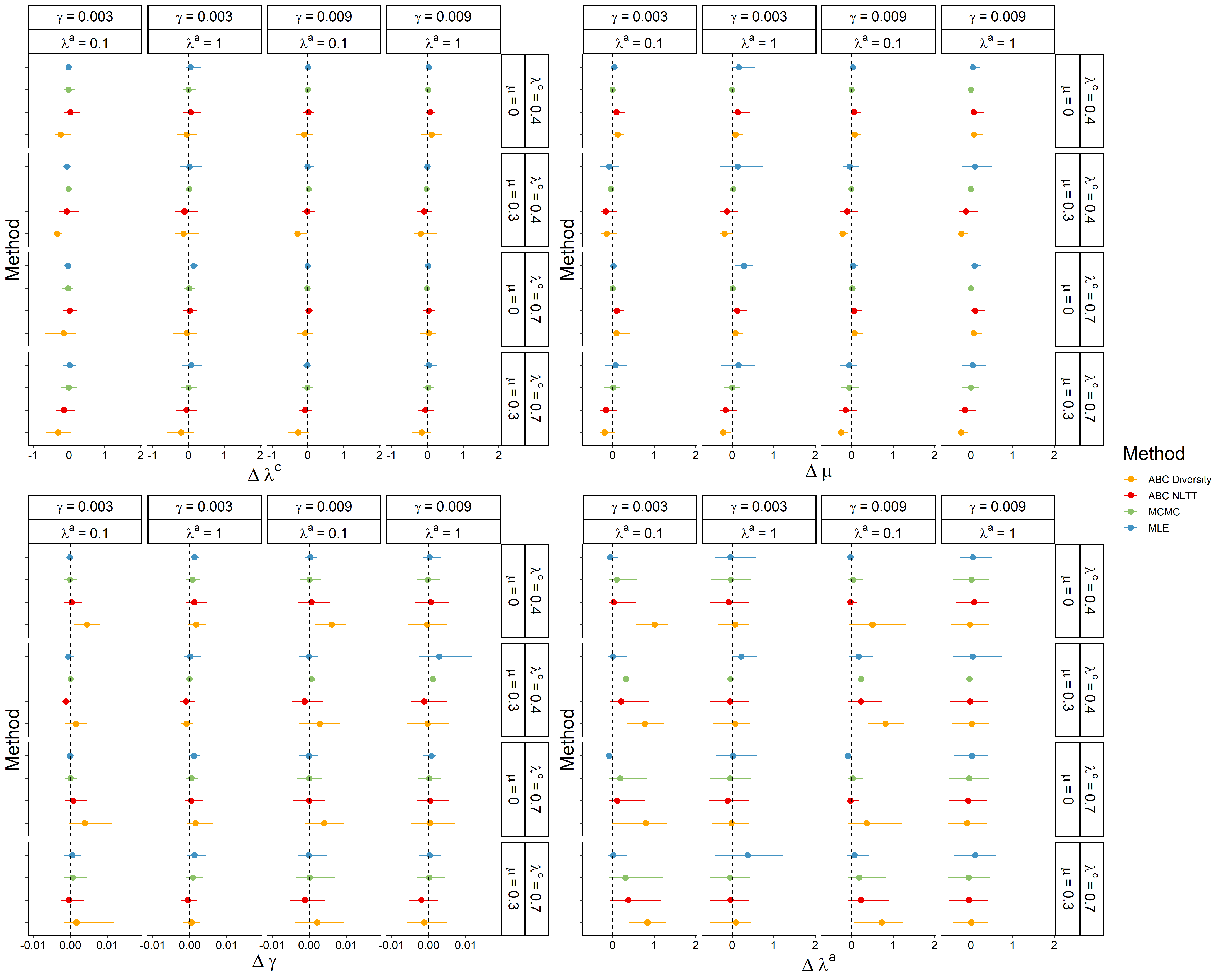

Figure S2. Results of inference of the **cladogenesis, extinction, colonization and anagenesis rates** using MLE, MCMC and ABC. Lines show 95% confidence intervals, dots the median value. The ABC results are from the analyses using the **narrower prior scale** (See Table 2). Plots show the Δ-net diversification rate, that is, the difference between the estimated value and the true value used for generating the observed simulations. The true values are shown at the top of each column and to the right of each row. The distribution for each method combines the 10 replicates for each parameter set. For MLE, these replicates result in 10 data points, whereas for MCMC and ABC these replicates result in 10 posterior distributions combined. In the case of ABC-SMC, the posterior distribution for each replicate is based on 500 particles from the last iteration step of the algorithm. ABC All - all statistics; ABC Diversity - three diversity-related statistics; ABC NLTT - three NLTT statistics; ABC Phylogenetic - five phylogenetic statistics; MLE - Maximum likelihood estimation; MCMC - Markov Chain Monte Carlo; γ - colonization rate; μ - extinction rate; λ^c^ – cladogenesis rate; λ^a^ – anagenesis rate.

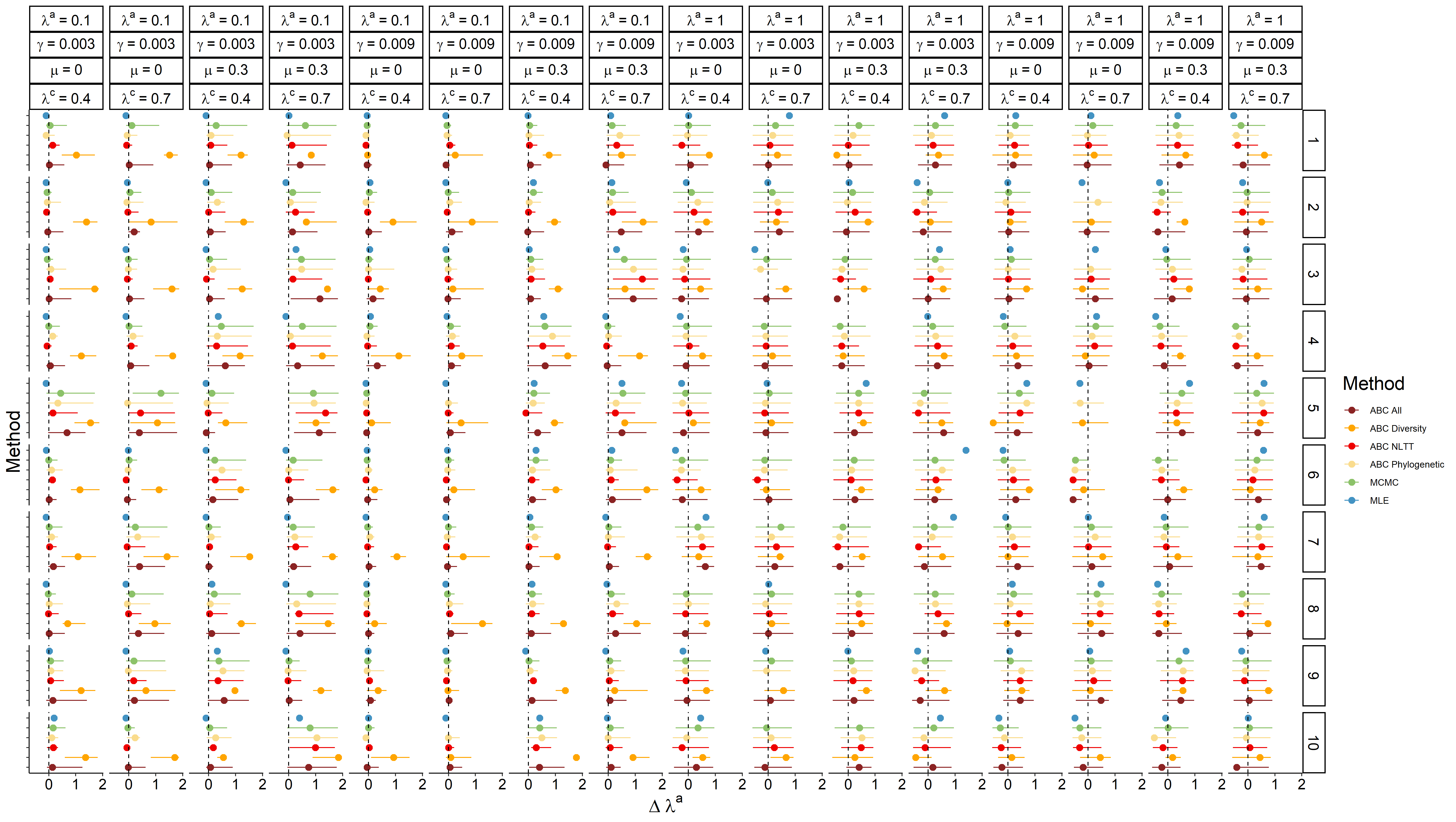

Figure S3. Results of inference of the **anagenesis rate** using MLE, MCMC and ABC for each individual observed dataset (a single replicate) (95% confidence interval). The ABC results are from the analyses using the **broader prior scale** (Table 2). Plots show the Δ- anagenesis, that is, the difference between the estimated values and the true values used for generating the observed simulations. The true values are shown at the top of each column. The numbers to the right of each row indicate the replicate. In the ABC-SMC algorithm, 500 particles in the last iteration are used as posterior for each replicate. For MLE these are therefore 10 data points, whereas for MCMC and ABC these are 10 posterior distributions combined. ABC All - all statistics; ABC Diversity - three diversity-related statistics; ABC NLTT - three NLTT statistics; ABC Phylogenetic - five phylogenetic statistics; MLE - Maximum likelihood estimation; MCMC - Markov Chain Monte Carlo; γ - colonization rate; μ - extinction rate; λ^c^ – cladogenesis rate; λ^a^ – anagenesis rate.

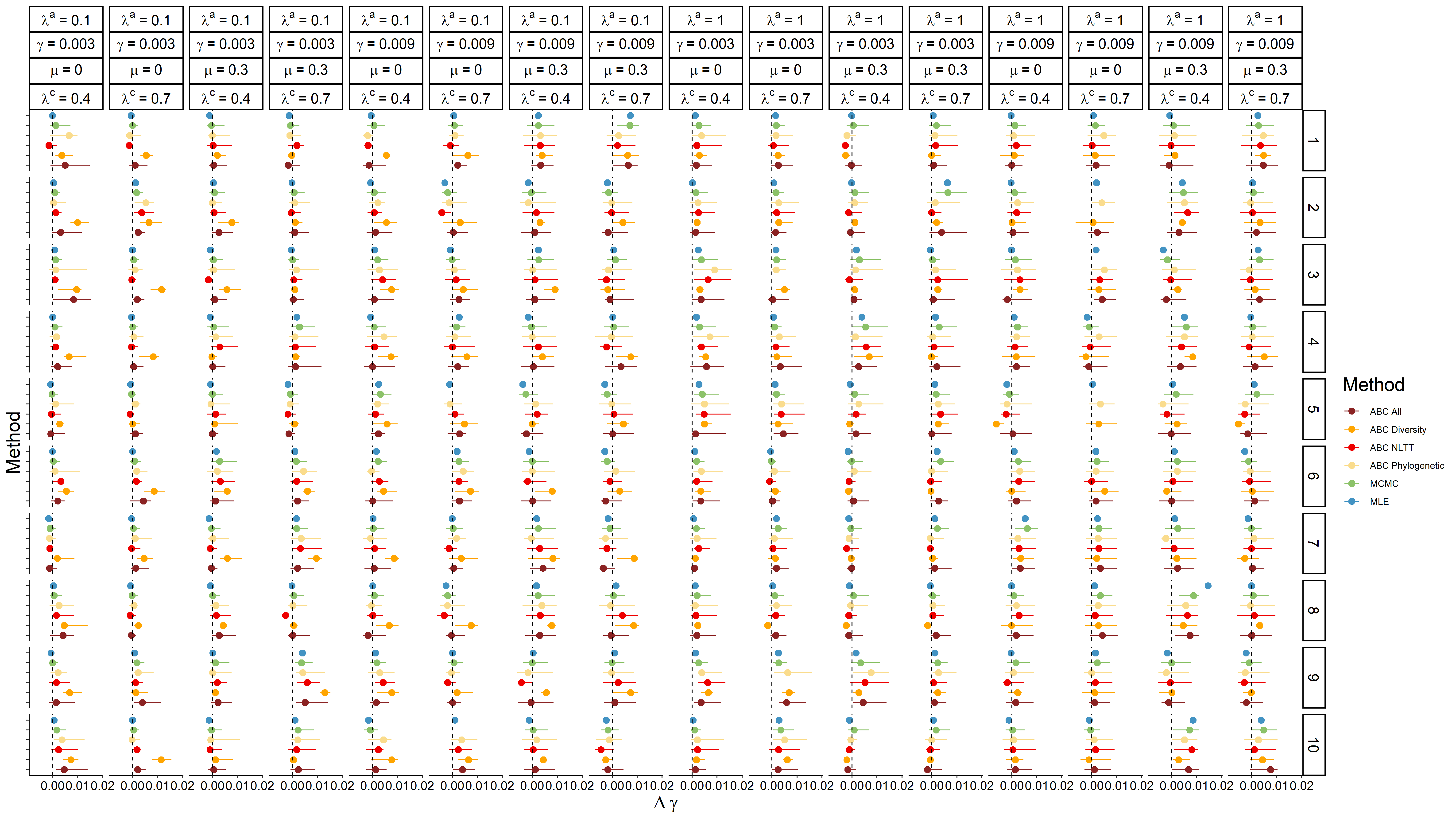

Figure S4. Results of inference of the **colonization rate** using MLE, MCMC and ABC for each individual observed dataset (a single replicate) (95% confidence interval). The ABC results are from the analyses using the **broader prior scale** (Table 2). Plots show the Δ- colonization, that is, the difference between the estimated values and the true values used for generating the observed simulations. The true values are shown at the top of each column. The numbers to the right of each row indicate the replicate. In the ABC-SMC algorithm, 500 particles in the last iteration are used as posterior for each replicate. For MLE these are therefore 10 data points, whereas for MCMC and ABC these are 10 posterior distributions combined. ABC All - all statistics; ABC Diversity - three diversity-related statistics; ABC NLTT - three NLTT statistics; ABC Phylogenetic - five phylogenetic statistics; MLE - Maximum likelihood estimation; MCMC - Markov Chain Monte Carlo; γ - colonization rate; μ - extinction rate; λ^c^ – cladogenesis rate; λ^a^ – anagenesis rate.

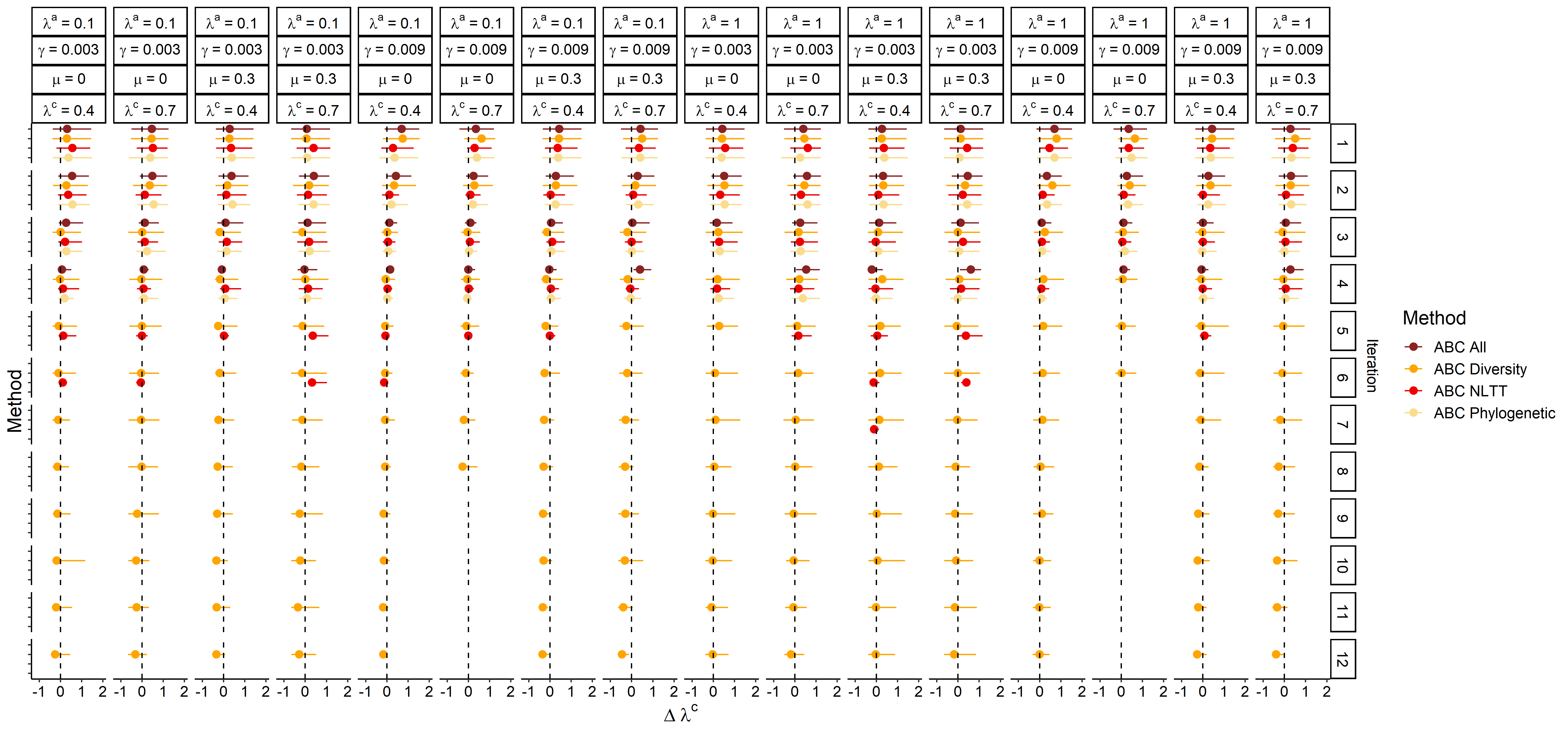

Figure S5. Results of inference of the **cladogenesis rate** **through increasing iterations** using ABC method with different groups of summary statistics. The ABC results are from the analyses using the **broader prior scale** (Table 2). Plots show the Δ- cladogenesis, that is, the difference between the estimated values and the true values used for generating the observed simulations. The true values are shown at the top of each column. The right of each row shows the number of the iteration. The distribution for each method combines the replicates for each parameter set at a specific iteration. ABC All - all statistics; ABC Diversity - three diversity-related statistics; ABC NLTT - three NLTT statistics; ABC Phylogenetic - five phylogenetic statistics; γ - colonization rate; μ - extinction rate; λ^c^ – cladogenesis rate; λ^a^ – anagenesis rate.

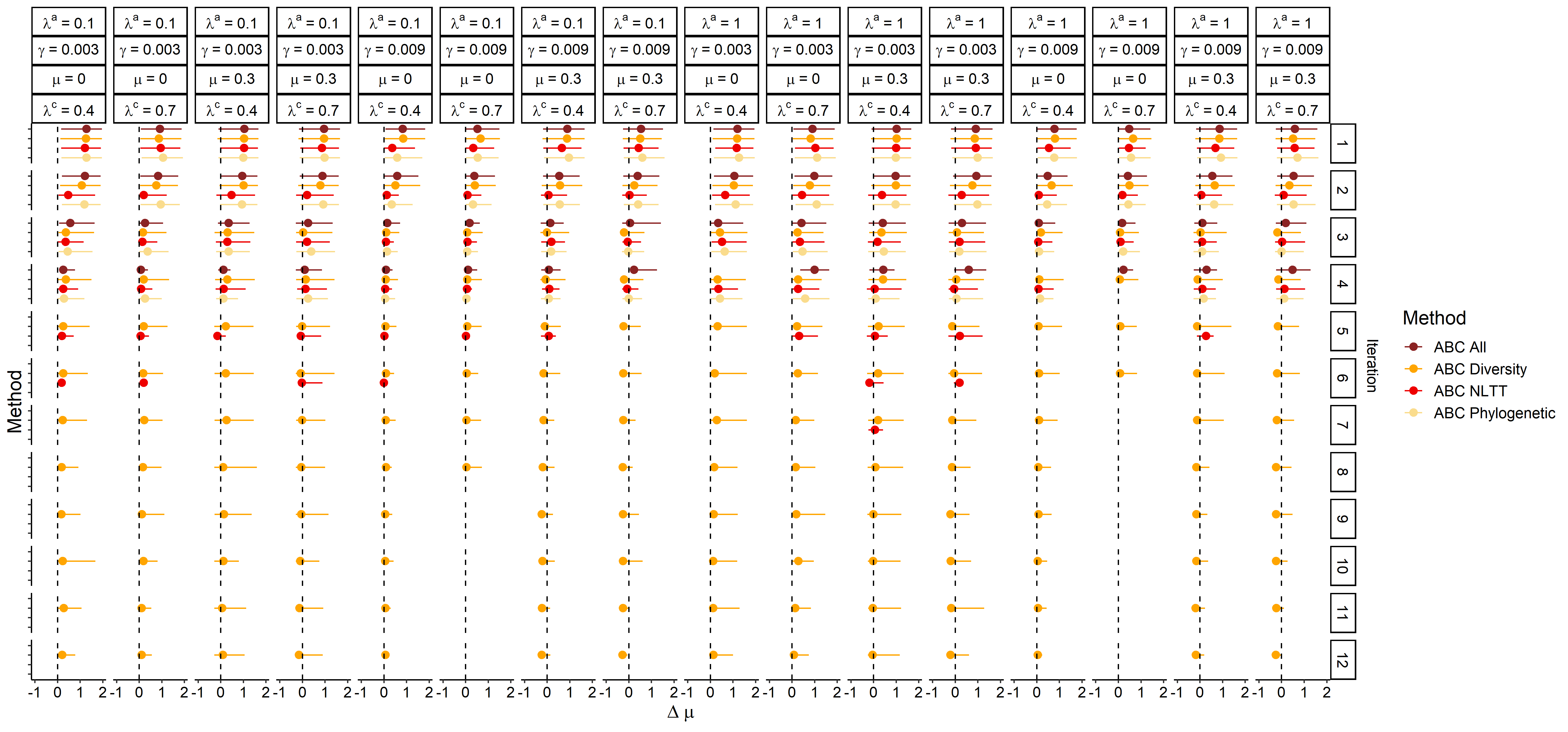

Figure S6. Results of inference of the **extinction rate** **through increasing iterations** using ABC method with different groups of summary statistics. The ABC results are from the analyses using the **broader prior scale** (Table 2). Plots show the Δ- extinction, that is, the difference between the estimated values and the true values used for generating the observed simulations. The true values are shown at the top of each column. The right of each row shows the number of the iteration. The distribution for each method combines the replicates for each parameter set at a specific iteration. ABC All - all statistics; ABC Diversity - three diversity-related statistics; ABC NLTT - three NLTT statistics; ABC Phylogenetic - five phylogenetic statistics; γ - colonization rate; μ - extinction rate; λ^c^ – cladogenesis rate; λ^a^ – anagenesis rate.

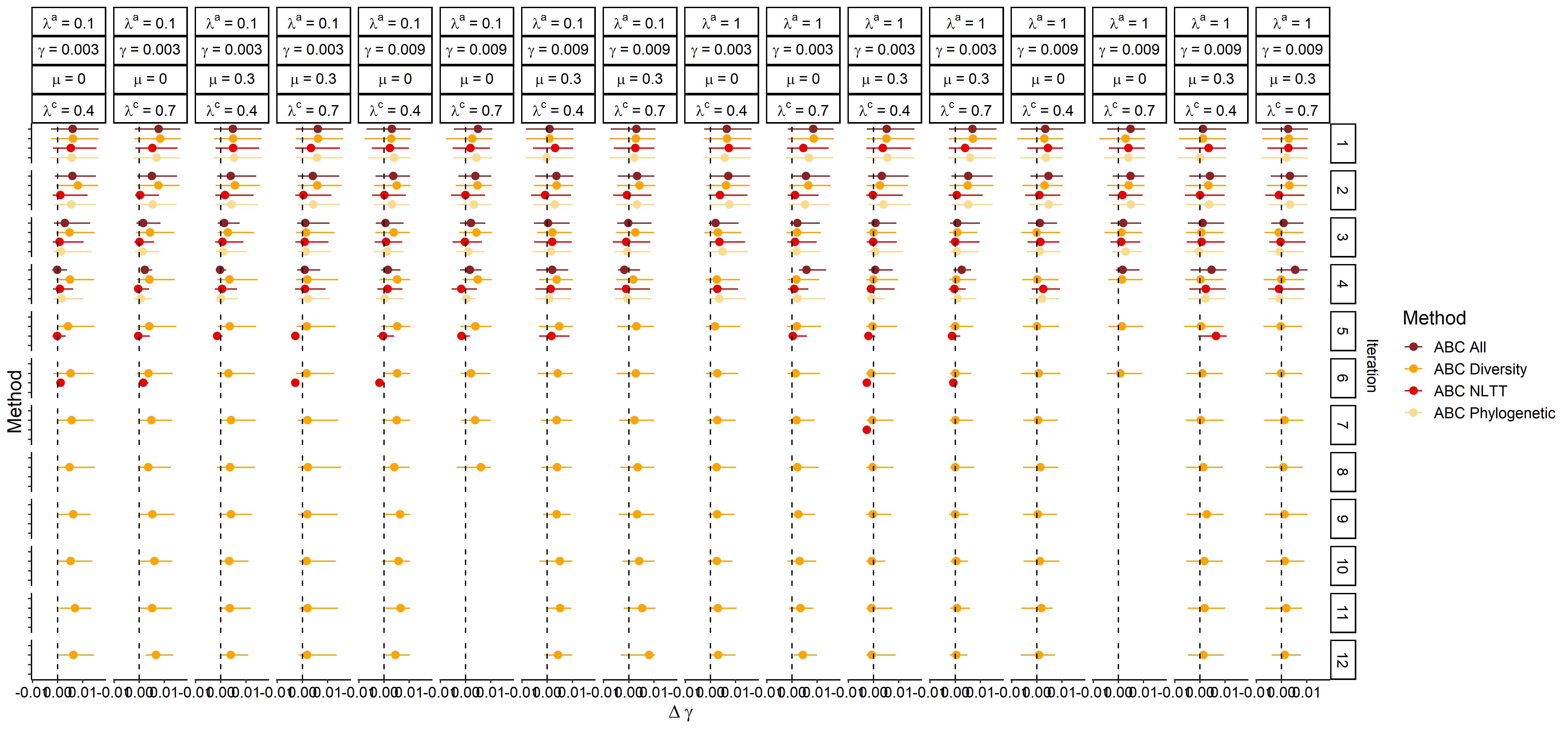

Figure S7. Results of inference of the **colonization rate** **through increasing iterations** using ABC method with different groups of summary statistics. The ABC results are from the analyses using the **broader prior scale** (Table 2). Plots show the Δ- colonization, that is, the difference between the estimated values and the true values used for generating the observed simulations. The true values are shown at the top of each column. The right of each row shows the number of the iteration. The distribution for each method combines the replicates for each parameter set at a specific iteration. ABC All - all statistics; ABC Diversity - three diversity-related statistics; ABC NLTT - three NLTT statistics; ABC Phylogenetic - five phylogenetic statistics; γ - colonization rate; μ - extinction rate; λ^c^ – cladogenesis rate; λ^a^ – anagenesis rate.

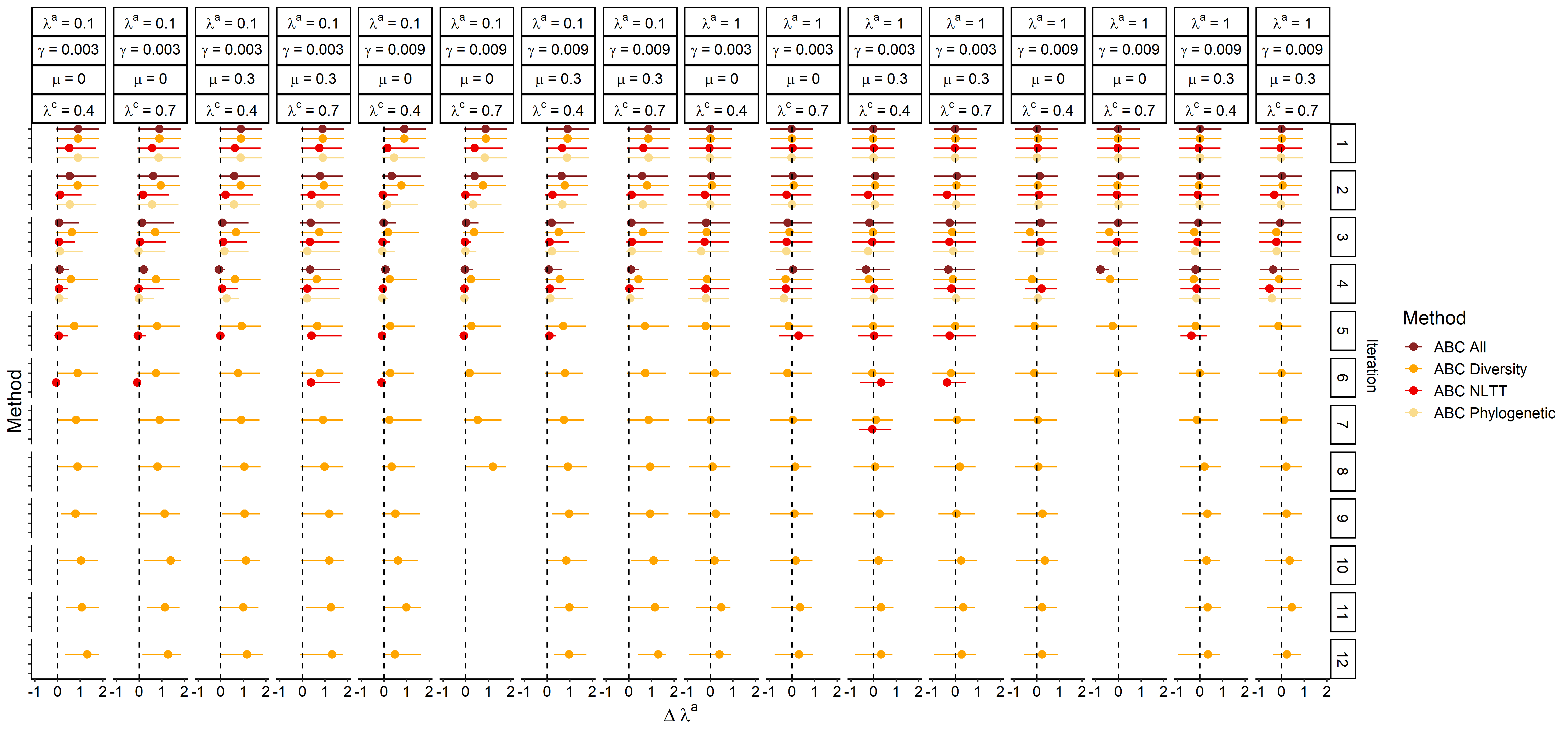

Figure S8. Results of inference of the **anagenesis rate** **through increasing iterations** using ABC method with different groups of summary statistics. The ABC results are from the analyses using the broader prior scale (Table 2). Plots show the Δ- anagenesis, that is, the difference between the estimated values and the true values used for generating the observed simulations. The true values are shown at the top of each column. The right of each row shows the number of the iteration. The distribution for each method combines the replicates for each parameter set at a specific iteration. ABC All - all statistics; ABC Diversity - three diversity-related statistics; ABC NLTT - three NLTT statistics; ABC Phylogenetic - five phylogenetic statistics; γ - colonization rate; μ - extinction rate; λ^c^ – cladogenesis rate; λ^a^ – anagenesis rate.

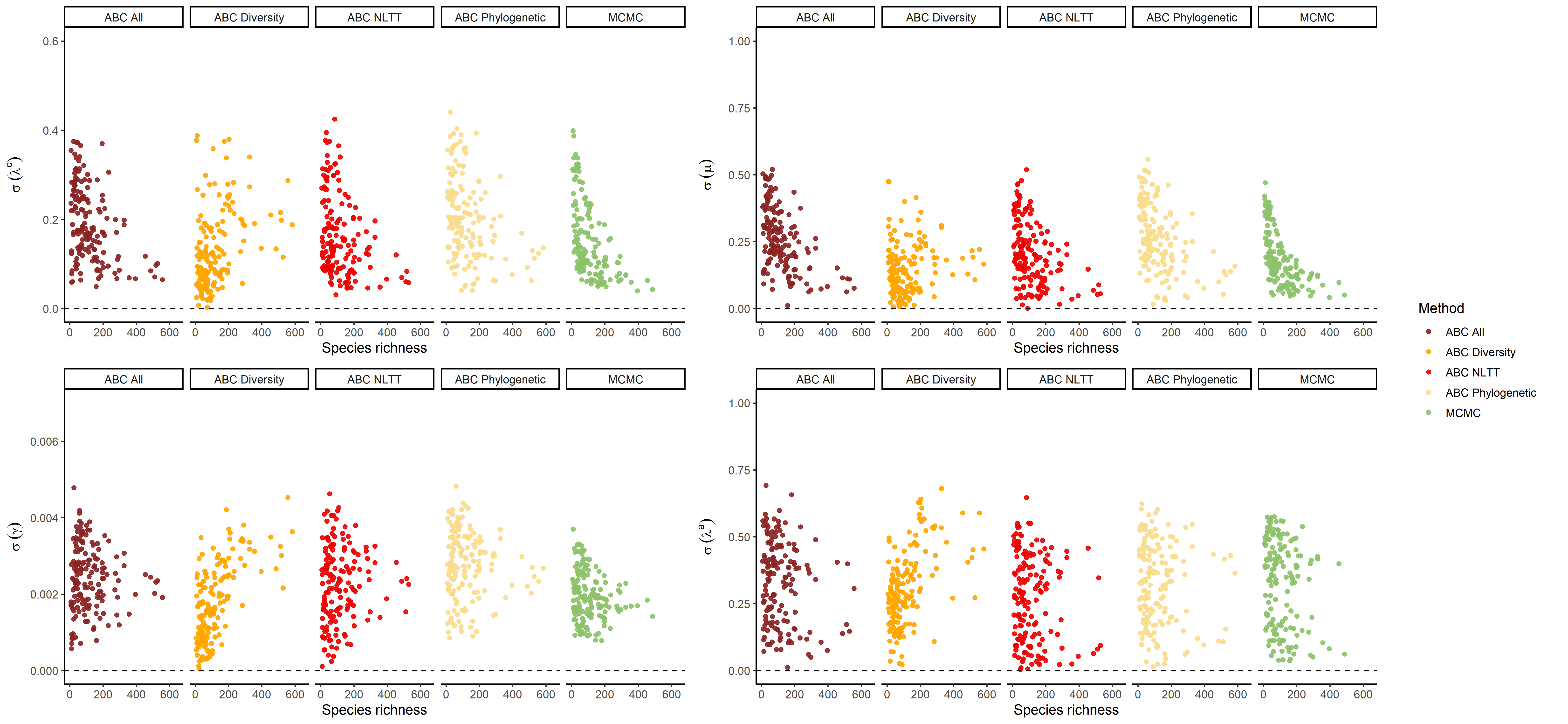

Figure S9. The relationship between the **species richness** and the relative difference in estimating the different rates. Different colors indicate different estimation methods. The points show the **standard deviation** of all the accepted particles in each posterior distribution using MCMC and ABC.

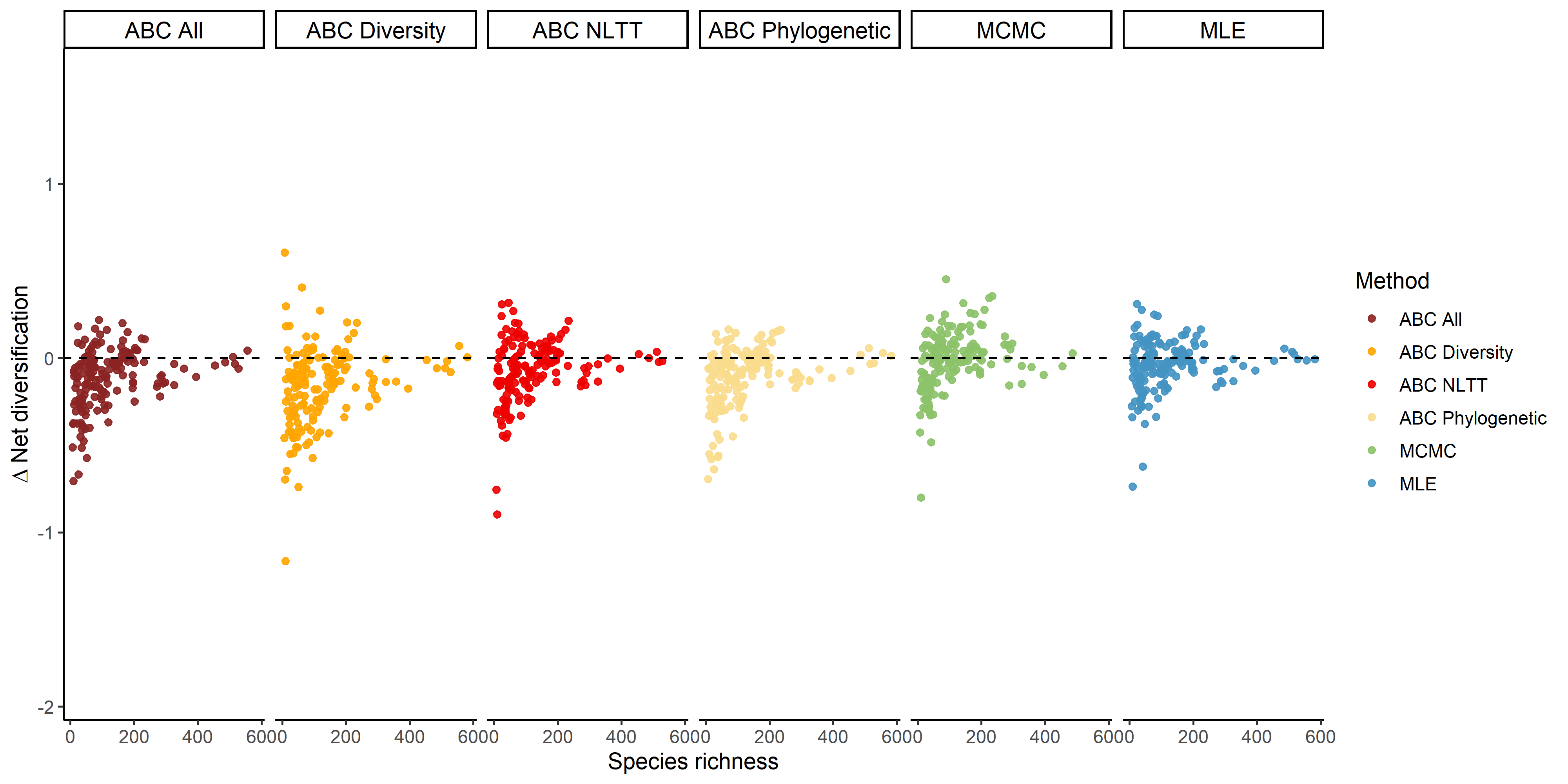

Figure S10. Relative difference in the estimates of **net diversification rate** (estimated value minus true value) against the **total species richness** per simulated dataset. Colors indicate different estimation methods. The points show the point estimation of MLE or the **median** value of the posterior distribution using MCMC and ABC.

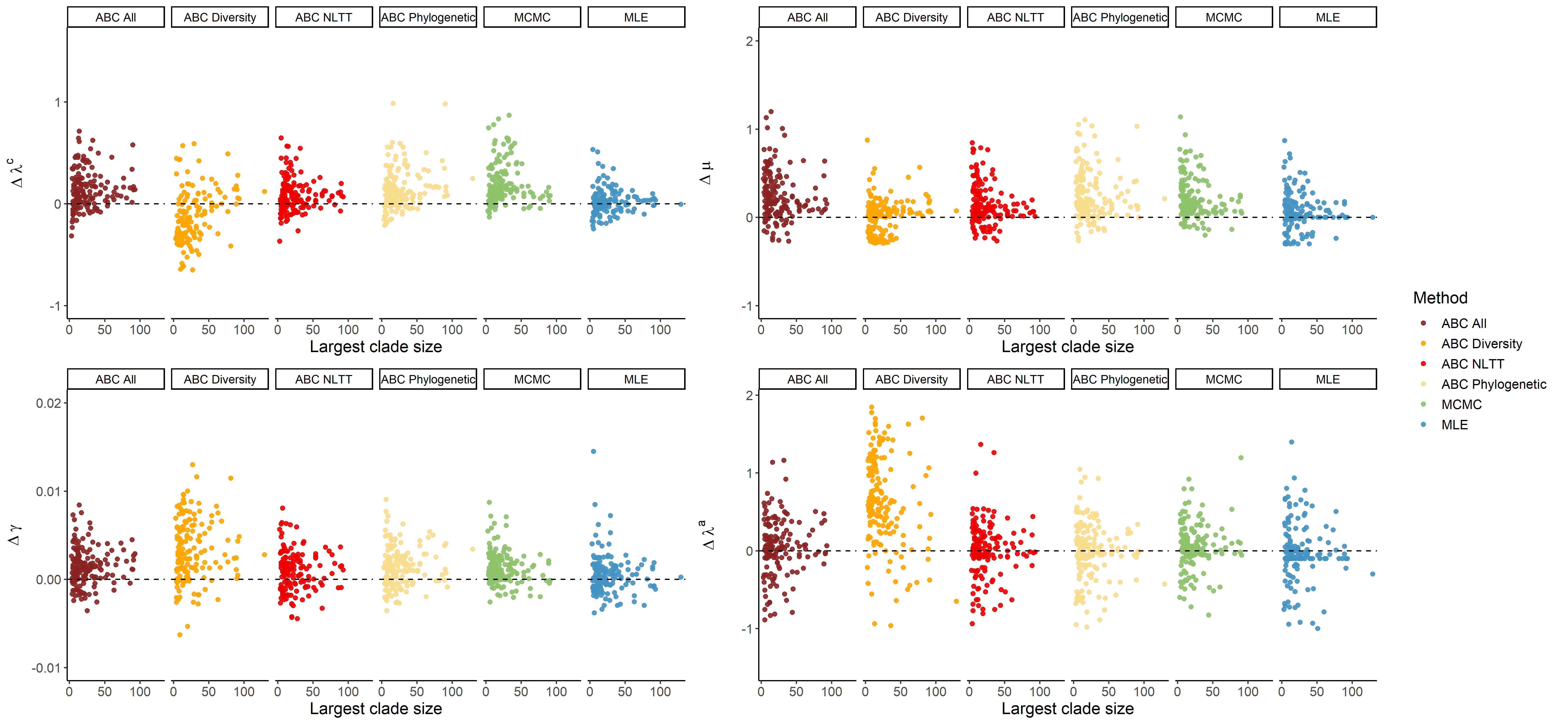

Figure S11. The relative difference in estimating rates for each dataset, plotted against the **size of the largest clade** in a dataset. Colors indicate different estimation methods. The points show the point estimation of MLE or the **median** value of the posterior distribution using MCMC and ABC.

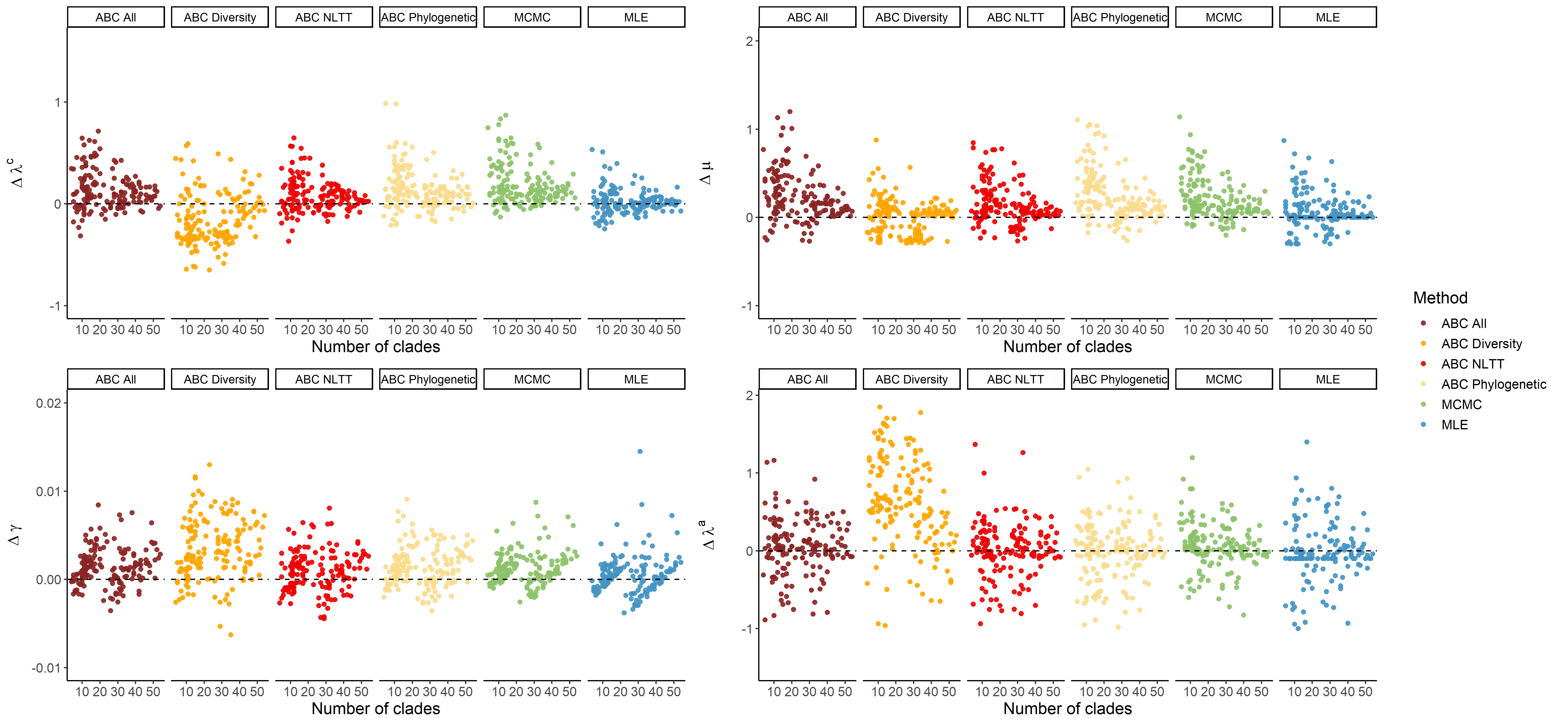

Figure S12. The relationship between the **number of clades** of the whole island community in a simulated dataset and the relative difference in estimating parameters. Colors indicate different estimation methods. The points show the point estimation of MLE or the **median** value of the posterior distribution using MCMC and ABC.

Table S1. Summary statistics of each of the 160 observed datasets. Column 2-5 are the generating parameter values for each observed dataset, and columns 6-14 are the calculated summary statistics. γ - colonization rate; μ - extinction rate; λ^c^ - cladogenesis rate; λ^a^ - anagenesis rate; NLTT_total_ - number of island lineages through time; NLTT_singleton-end_ - number of singleton island endemics through time; NLTT_non-end_ - number of non-endemic island species through time; N_total_ - total number of extant species on the island; N_end_ - number of island endemic species; N_non-end_ - number of non-endemic island species; SD-CT - standard deviation of colonization time among island clades; SD-CS - standard deviation of clade size; Largest clade - number of species of the largest clade.

| Dataset | λ^c^ | μ | γ | λ^a^ | NLTT_total_ | NLTT_singleton-end_ | NLTT_non-end_ | SD-CT | *N*_total_ | *N*_end_ | *N*_non-end_ | SD-CS | Largest clade |
| --- | --- | --- | --- | --- | --- | --- | --- | --- | --- | --- | --- | --- | --- |
| 1 | 0.4 | 0 | 0.003 | 0.1 | 37.7 | 0.0 | 4.6 | 1.2 | 25 | 20 | 5 | 1.4 | 7 |
| 2 | 0.4 | 0 | 0.003 | 0.1 | 82.0 | 0.0 | 17.8 | 1.4 | 48 | 41 | 7 | 4.2 | 20 |
| 3 | 0.4 | 0 | 0.003 | 0.1 | 117.2 | 0.0 | 13.8 | 1.0 | 60 | 54 | 6 | 3.2 | 14 |
| 4 | 0.4 | 0 | 0.003 | 0.1 | 96.2 | 0.0 | 7.3 | 1.6 | 66 | 61 | 5 | 4.4 | 16 |
| 5 | 0.4 | 0 | 0.003 | 0.1 | 108.3 | 0.0 | 1.1 | 1.8 | 56 | 54 | 2 | 4.0 | 15 |
| 6 | 0.4 | 0 | 0.003 | 0.1 | 69.9 | 4.2 | 9.8 | 1.4 | 36 | 31 | 5 | 2.3 | 9 |
| 7 | 0.4 | 0 | 0.003 | 0.1 | 35.3 | 0.0 | 6.0 | 1.6 | 20 | 17 | 3 | 3.2 | 11 |
| 8 | 0.4 | 0 | 0.003 | 0.1 | 59.8 | 3.2 | 13.1 | 1.5 | 32 | 24 | 8 | 2.2 | 9 |
| 9 | 0.4 | 0 | 0.003 | 0.1 | 100.9 | 2.3 | 6.9 | 1.6 | 53 | 48 | 5 | 7.0 | 23 |
| 10 | 0.4 | 0 | 0.003 | 0.1 | 79.8 | 8.6 | 9.7 | 1.1 | 46 | 41 | 5 | 2.0 | 8 |
| 11 | 0.7 | 0 | 0.003 | 0.1 | 89.2 | 0.0 | 2.8 | 1.2 | 76 | 73 | 3 | 8.7 | 27 |
| 12 | 0.7 | 0 | 0.003 | 0.1 | 295.4 | 2.8 | 7.9 | 1.4 | 231 | 226 | 5 | 17.6 | 68 |
| 13 | 0.7 | 0 | 0.003 | 0.1 | 130.3 | 0.0 | 8.8 | 1.7 | 95 | 89 | 6 | 9.5 | 33 |
| 14 | 0.7 | 0 | 0.003 | 0.1 | 151.8 | 0.0 | 6.6 | 1.5 | 107 | 104 | 3 | 15.8 | 61 |
| 15 | 0.7 | 0 | 0.003 | 0.1 | 248.7 | 0.9 | 0.0 | 1.6 | 181 | 180 | 1 | 26.4 | 90 |
| 16 | 0.7 | 0 | 0.003 | 0.1 | 69.6 | 0.5 | 11.8 | 1.4 | 51 | 43 | 8 | 4.0 | 13 |
| 17 | 0.7 | 0 | 0.003 | 0.1 | 132.8 | 0.0 | 2.0 | 1.4 | 96 | 93 | 3 | 7.5 | 24 |
| 18 | 0.7 | 0 | 0.003 | 0.1 | 206.2 | 0.0 | 1.9 | 1.5 | 160 | 158 | 2 | 25.8 | 86 |
| 19 | 0.7 | 0 | 0.003 | 0.1 | 243.0 | 0.0 | 2.3 | 1.5 | 197 | 195 | 2 | 11.4 | 41 |
| 20 | 0.7 | 0 | 0.003 | 0.1 | 178.9 | 0.0 | 12.1 | 1.5 | 146 | 139 | 7 | 20.0 | 81 |
| 21 | 0.4 | 0.3 | 0.003 | 0.1 | 13.9 | 2.9 | 2.3 | 1.2 | 9 | 6 | 3 | 1.8 | 6 |
| 22 | 0.4 | 0.3 | 0.003 | 0.1 | 50.3 | 4.3 | 4.3 | 1.7 | 31 | 26 | 5 | 3.4 | 14 |
| 23 | 0.4 | 0.3 | 0.003 | 0.1 | 44.5 | 0.0 | 4.9 | 1.9 | 24 | 19 | 5 | 1.5 | 7 |
| 24 | 0.4 | 0.3 | 0.003 | 0.1 | 11.5 | 3.6 | 1.9 | 1.2 | 7 | 5 | 2 | 0.9 | 4 |
| 25 | 0.4 | 0.3 | 0.003 | 0.1 | 22.8 | 4.7 | 3.7 | 1.9 | 12 | 8 | 4 | 1.3 | 5 |
| 26 | 0.4 | 0.3 | 0.003 | 0.1 | 35.2 | 0.0 | 2.2 | 1.7 | 23 | 19 | 4 | 0.9 | 5 |
| 27 | 0.4 | 0.3 | 0.003 | 0.1 | 26.7 | 0.0 | 6.6 | 1.5 | 14 | 10 | 4 | 1.0 | 5 |
| 28 | 0.4 | 0.3 | 0.003 | 0.1 | 71.5 | 3.4 | 3.0 | 1.8 | 38 | 33 | 5 | 3.9 | 12 |
| 29 | 0.4 | 0.3 | 0.003 | 0.1 | 37.5 | 13.4 | 2.6 | 1.6 | 20 | 17 | 3 | 1.3 | 5 |
| 30 | 0.4 | 0.3 | 0.003 | 0.1 | 24.6 | 0.0 | 4.7 | 1.6 | 16 | 11 | 5 | 2.2 | 7 |
| 31 | 0.7 | 0.3 | 0.003 | 0.1 | 32.0 | 2.5 | 0.7 | 1.4 | 24 | 23 | 1 | 5.8 | 16 |
| 32 | 0.7 | 0.3 | 0.003 | 0.1 | 46.9 | 0.0 | 2.9 | 1.5 | 32 | 29 | 3 | 3.6 | 14 |
| 33 | 0.7 | 0.3 | 0.003 | 0.1 | 102.3 | 5.4 | 1.0 | 0.9 | 89 | 88 | 1 | 10.8 | 32 |
| 34 | 0.7 | 0.3 | 0.003 | 0.1 | 49.2 | 3.6 | 0.8 | 1.4 | 39 | 37 | 2 | 3.1 | 11 |
| 35 | 0.7 | 0.3 | 0.003 | 0.1 | 57.4 | 1.9 | 0.0 | 1.6 | 35 | 34 | 1 | 5.4 | 16 |
| 36 | 0.7 | 0.3 | 0.003 | 0.1 | 68.1 | 3.5 | 2.6 | 1.6 | 48 | 44 | 4 | 4.3 | 14 |
| 37 | 0.7 | 0.3 | 0.003 | 0.1 | 73.6 | 8.8 | 4.5 | 1.8 | 44 | 38 | 6 | 3.7 | 14 |
| 38 | 0.7 | 0.3 | 0.003 | 0.1 | 55.0 | 0.0 | 0.3 | 1.3 | 46 | 45 | 1 | 5.2 | 20 |
| 39 | 0.7 | 0.3 | 0.003 | 0.1 | 105.4 | 7.5 | 9.6 | 1.4 | 79 | 68 | 11 | 5.9 | 27 |
| 40 | 0.7 | 0.3 | 0.003 | 0.1 | 52.4 | 9.9 | 0.4 | 1.4 | 38 | 37 | 1 | 2.5 | 9 |
| 41 | 0.4 | 0 | 0.009 | 0.1 | 132.0 | 1.7 | 41.2 | 1.3 | 91 | 69 | 22 | 2.3 | 14 |
| 42 | 0.4 | 0 | 0.009 | 0.1 | 202.8 | 11.2 | 12.4 | 1.4 | 113 | 103 | 10 | 2.1 | 10 |
| 43 | 0.4 | 0 | 0.009 | 0.1 | 268.9 | 16.5 | 31.0 | 1.6 | 154 | 135 | 19 | 4.3 | 23 |
| 44 | 0.4 | 0 | 0.009 | 0.1 | 224.0 | 11.6 | 13.4 | 1.7 | 121 | 109 | 12 | 3.0 | 13 |
| 45 | 0.4 | 0 | 0.009 | 0.1 | 259.2 | 3.4 | 32.8 | 1.4 | 158 | 139 | 19 | 2.3 | 12 |
| 46 | 0.4 | 0 | 0.009 | 0.1 | 261.9 | 4.0 | 36.0 | 1.5 | 161 | 138 | 23 | 5.0 | 35 |
| 47 | 0.4 | 0 | 0.009 | 0.1 | 192.7 | 6.9 | 18.3 | 1.4 | 111 | 99 | 12 | 2.7 | 15 |
| 48 | 0.4 | 0 | 0.009 | 0.1 | 186.0 | 4.4 | 36.2 | 1.4 | 117 | 97 | 20 | 3.7 | 22 |
| 49 | 0.4 | 0 | 0.009 | 0.1 | 322.9 | 12.6 | 31.0 | 1.5 | 182 | 161 | 21 | 6.4 | 37 |
| 50 | 0.4 | 0 | 0.009 | 0.1 | 301.1 | 5.2 | 15.8 | 1.5 | 167 | 156 | 11 | 4.8 | 18 |
| 51 | 0.7 | 0 | 0.009 | 0.1 | 381.2 | 2.1 | 21.5 | 1.4 | 297 | 283 | 14 | 8.4 | 40 |
| 52 | 0.7 | 0 | 0.009 | 0.1 | 267.1 | 0.0 | 6.3 | 1.4 | 201 | 194 | 7 | 5.7 | 20 |
| 53 | 0.7 | 0 | 0.009 | 0.1 | 592.3 | 0.0 | 15.2 | 1.5 | 486 | 475 | 11 | 19.3 | 91 |
| 54 | 0.7 | 0 | 0.009 | 0.1 | 352.9 | 8.3 | 10.0 | 1.3 | 272 | 260 | 12 | 8.2 | 51 |
| 55 | 0.7 | 0 | 0.009 | 0.1 | 641.3 | 0.0 | 10.0 | 1.4 | 511 | 501 | 10 | 20.7 | 93 |
| 56 | 0.7 | 0 | 0.009 | 0.1 | 541.4 | 1.1 | 15.4 | 1.5 | 395 | 380 | 15 | 10.0 | 54 |
| 57 | 0.7 | 0 | 0.009 | 0.1 | 444.0 | 1.7 | 10.7 | 1.4 | 357 | 348 | 9 | 10.8 | 54 |
| 58 | 0.7 | 0 | 0.009 | 0.1 | 251.7 | 0.0 | 9.9 | 1.4 | 195 | 186 | 9 | 11.4 | 63 |
| 59 | 0.7 | 0 | 0.009 | 0.1 | 379.4 | 0.0 | 15.1 | 1.5 | 283 | 268 | 15 | 8.9 | 44 |
| 60 | 0.7 | 0 | 0.009 | 0.1 | 687.8 | 0.0 | 9.9 | 1.7 | 528 | 513 | 15 | 14.7 | 54 |
| 61 | 0.4 | 0.3 | 0.009 | 0.1 | 107.2 | 19.3 | 17.7 | 1.5 | 65 | 53 | 12 | 1.5 | 9 |
| 62 | 0.4 | 0.3 | 0.009 | 0.1 | 95.7 | 12.5 | 13.7 | 1.4 | 59 | 49 | 10 | 2.1 | 10 |
| 63 | 0.4 | 0.3 | 0.009 | 0.1 | 140.7 | 10.0 | 10.1 | 1.4 | 93 | 79 | 14 | 2.7 | 14 |
| 64 | 0.4 | 0.3 | 0.009 | 0.1 | 117.6 | 21.2 | 3.4 | 1.5 | 68 | 61 | 7 | 2.4 | 11 |
| 65 | 0.4 | 0.3 | 0.009 | 0.1 | 88.0 | 17.5 | 6.2 | 1.6 | 45 | 37 | 8 | 1.4 | 6 |
| 66 | 0.4 | 0.3 | 0.009 | 0.1 | 103.3 | 22.1 | 11.5 | 1.6 | 58 | 47 | 11 | 1.9 | 10 |
| 67 | 0.4 | 0.3 | 0.009 | 0.1 | 55.3 | 14.5 | 10.0 | 1.2 | 35 | 24 | 11 | 0.8 | 5 |
| 68 | 0.4 | 0.3 | 0.009 | 0.1 | 114.1 | 16.3 | 14.5 | 1.3 | 72 | 60 | 12 | 2.0 | 12 |
| 69 | 0.4 | 0.3 | 0.009 | 0.1 | 99.1 | 6.0 | 10.1 | 1.4 | 71 | 59 | 12 | 3.6 | 20 |
| 70 | 0.4 | 0.3 | 0.009 | 0.1 | 134.3 | 19.7 | 6.7 | 1.3 | 73 | 66 | 7 | 1.5 | 9 |
| 71 | 0.7 | 0.3 | 0.009 | 0.1 | 277.5 | 14.5 | 7.9 | 1.5 | 202 | 188 | 14 | 4.8 | 29 |
| 72 | 0.7 | 0.3 | 0.009 | 0.1 | 226.5 | 13.5 | 6.2 | 1.5 | 165 | 158 | 7 | 7.9 | 37 |
| 73 | 0.7 | 0.3 | 0.009 | 0.1 | 309.2 | 9.0 | 0.6 | 1.7 | 234 | 230 | 4 | 9.7 | 35 |
| 74 | 0.7 | 0.3 | 0.009 | 0.1 | 120.1 | 10.8 | 11.8 | 1.5 | 97 | 85 | 12 | 2.8 | 12 |
| 75 | 0.7 | 0.3 | 0.009 | 0.1 | 255.6 | 12.3 | 4.7 | 1.4 | 174 | 169 | 5 | 14.4 | 77 |
| 76 | 0.7 | 0.3 | 0.009 | 0.1 | 265.6 | 7.7 | 9.8 | 1.5 | 179 | 173 | 6 | 8.4 | 39 |
| 77 | 0.7 | 0.3 | 0.009 | 0.1 | 227.2 | 1.7 | 6.1 | 1.8 | 163 | 154 | 9 | 6.6 | 25 |
| 78 | 0.7 | 0.3 | 0.009 | 0.1 | 179.1 | 15.0 | 8.1 | 1.7 | 132 | 120 | 12 | 7.0 | 36 |
| 79 | 0.7 | 0.3 | 0.009 | 0.1 | 171.4 | 8.2 | 10.1 | 1.4 | 134 | 122 | 12 | 6.0 | 29 |
| 80 | 0.7 | 0.3 | 0.009 | 0.1 | 161.0 | 1.6 | 6.6 | 1.3 | 114 | 110 | 4 | 4.9 | 28 |
| 81 | 0.4 | 0 | 0.003 | 1 | 84.8 | 16.9 | 4.7 | 1.3 | 55 | 53 | 2 | 3.1 | 12 |
| 82 | 0.4 | 0 | 0.003 | 1 | 61.1 | 14.9 | 0.6 | 1.7 | 33 | 31 | 2 | 2.5 | 11 |
| 83 | 0.4 | 0 | 0.003 | 1 | 59.6 | 10.6 | 2.5 | 1.2 | 37 | 34 | 3 | 1.5 | 6 |
| 84 | 0.4 | 0 | 0.003 | 1 | 51.6 | 7.5 | 1.8 | 1.2 | 35 | 31 | 4 | 2.2 | 9 |
| 85 | 0.4 | 0 | 0.003 | 1 | 70.1 | 5.4 | 2.7 | 1.4 | 39 | 34 | 5 | 0.9 | 5 |
| 86 | 0.4 | 0 | 0.003 | 1 | 126.4 | 8.0 | 2.8 | 1.4 | 70 | 67 | 3 | 2.9 | 13 |
| 87 | 0.4 | 0 | 0.003 | 1 | 118.1 | 19.1 | 0.5 | 1.4 | 65 | 64 | 1 | 6.0 | 27 |
| 88 | 0.4 | 0 | 0.003 | 1 | 51.1 | 6.4 | 1.2 | 1.5 | 38 | 36 | 2 | 3.5 | 11 |
| 89 | 0.4 | 0 | 0.003 | 1 | 122.9 | 21.1 | 2.3 | 1.5 | 67 | 63 | 4 | 4.1 | 18 |
| 90 | 0.4 | 0 | 0.003 | 1 | 130.8 | 4.7 | 1.4 | 1.6 | 85 | 83 | 2 | 8.2 | 32 |
| 91 | 0.7 | 0 | 0.003 | 1 | 115.5 | 4.2 | 0.3 | 1.7 | 107 | 105 | 2 | 9.4 | 33 |
| 92 | 0.7 | 0 | 0.003 | 1 | 154.3 | 8.4 | 0.3 | 1.4 | 117 | 115 | 2 | 9.1 | 32 |
| 93 | 0.7 | 0 | 0.003 | 1 | 109.1 | 2.7 | 1.6 | 1.5 | 90 | 87 | 3 | 9.8 | 37 |
| 94 | 0.7 | 0 | 0.003 | 1 | 240.9 | 2.8 | 1.3 | 1.5 | 195 | 193 | 2 | 15.7 | 60 |
| 95 | 0.7 | 0 | 0.003 | 1 | 245.1 | 7.1 | 2.1 | 1.3 | 196 | 194 | 2 | 20.8 | 89 |
| 96 | 0.7 | 0 | 0.003 | 1 | 190.3 | 0.0 | 1.2 | 1.3 | 143 | 142 | 1 | 14.8 | 51 |
| 97 | 0.7 | 0 | 0.003 | 1 | 136.8 | 5.7 | 0.1 | 1.6 | 111 | 110 | 1 | 8.7 | 30 |
| 98 | 0.7 | 0 | 0.003 | 1 | 119.4 | 4.4 | 1.5 | 1.6 | 104 | 103 | 1 | 9.1 | 36 |
| 99 | 0.7 | 0 | 0.003 | 1 | 141.0 | 5.3 | 0.9 | 1.4 | 119 | 115 | 4 | 7.5 | 30 |
| 100 | 0.7 | 0 | 0.003 | 1 | 109.1 | 6.5 | 0.6 | 1.7 | 84 | 80 | 4 | 6.5 | 26 |
| 101 | 0.4 | 0.3 | 0.003 | 1 | 8.9 | 4.6 | 0.0 | 1.9 | 8 | 8 | 0 | 1.2 | 4 |
| 102 | 0.4 | 0.3 | 0.003 | 1 | 37.7 | 8.1 | 0.9 | 1.6 | 23 | 22 | 1 | 2.1 | 8 |
| 103 | 0.4 | 0.3 | 0.003 | 1 | 20.8 | 0.5 | 1.8 | 1.2 | 17 | 15 | 2 | 0.8 | 4 |
| 104 | 0.4 | 0.3 | 0.003 | 1 | 12.4 | 8.7 | 2.7 | 1.4 | 10 | 5 | 5 | 0.3 | 3 |
| 105 | 0.4 | 0.3 | 0.003 | 1 | 32.9 | 14.9 | 0.9 | 1.2 | 16 | 15 | 1 | 0.9 | 5 |
| 106 | 0.4 | 0.3 | 0.003 | 1 | 37.6 | 5.8 | 0.0 | 0.8 | 23 | 23 | 0 | 2.2 | 8 |
| 107 | 0.4 | 0.3 | 0.003 | 1 | 17.8 | 1.1 | 1.8 | 1.6 | 11 | 10 | 1 | 1.3 | 5 |
| 108 | 0.4 | 0.3 | 0.003 | 1 | 19.0 | 8.1 | 0.0 | 1.5 | 12 | 12 | 0 | 2.0 | 7 |
| 109 | 0.4 | 0.3 | 0.003 | 1 | 31.2 | 14.5 | 1.1 | 1.1 | 21 | 19 | 2 | 1.3 | 6 |
| 110 | 0.4 | 0.3 | 0.003 | 1 | 26.3 | 12.4 | 0.0 | 1.6 | 12 | 12 | 0 | 0.8 | 4 |
| 111 | 0.7 | 0.3 | 0.003 | 1 | 42.3 | 6.3 | 0.4 | 1.5 | 30 | 29 | 1 | 3.1 | 11 |
| 112 | 0.7 | 0.3 | 0.003 | 1 | 59.9 | 1.1 | 1.4 | 1.0 | 60 | 58 | 2 | 2.9 | 12 |
| 113 | 0.7 | 0.3 | 0.003 | 1 | 82.6 | 5.9 | 1.0 | 1.6 | 54 | 52 | 2 | 6.2 | 23 |
| 114 | 0.7 | 0.3 | 0.003 | 1 | 40.9 | 8.5 | 0.2 | 1.3 | 33 | 32 | 1 | 1.9 | 7 |
| 115 | 0.7 | 0.3 | 0.003 | 1 | 37. 6 | 5.6 | 1.4 | 1.6 | 27 | 24 | 3 | 2.9 | 10 |
| 116 | 0.7 | 0.3 | 0.003 | 1 | 91.1 | 3.1 | 0.1 | 1.4 | 76 | 75 | 1 | 3.6 | 14 |
| 117 | 0.7 | 0.3 | 0.003 | 1 | 75. 3 | 0.9 | 0.1 | 1.4 | 65 | 64 | 1 | 4.7 | 18 |
| 118 | 0.7 | 0.3 | 0.003 | 1 | 100.5 | 13.2 | 0.0 | 1.4 | 62 | 62 | 0 | 7.9 | 29 |
| 119 | 0.7 | 0.3 | 0.003 | 1 | 136.2 | 7.4 | 1.7 | 1.5 | 96 | 94 | 2 | 7.1 | 22 |
| 120 | 0.7 | 0.3 | 0.003 | 1 | 50.8 | 0.7 | 0.3 | 1.5 | 45 | 44 | 1 | 4.2 | 13 |
| 121 | 0.4 | 0 | 0.009 | 1 | 279.8 | 26.9 | 3.3 | 1.3 | 166 | 162 | 4 | 4.9 | 20 |
| 122 | 0.4 | 0 | 0.009 | 1 | 241.7 | 30.8 | 3.9 | 1.4 | 136 | 130 | 6 | 3.8 | 22 |
| 123 | 0.4 | 0 | 0.009 | 1 | 128.1 | 20.0 | 2.9 | 1.5 | 77 | 70 | 7 | 1.7 | 8 |
| 124 | 0.4 | 0 | 0.009 | 1 | 322.7 | 26.4 | 3.6 | 1.4 | 187 | 182 | 5 | 4.4 | 20 |
| 125 | 0.4 | 0 | 0.009 | 1 | 219.1 | 12.5 | 1.1 | 1.4 | 119 | 117 | 2 | 2.3 | 9 |
| 126 | 0.4 | 0 | 0.009 | 1 | 299.2 | 33.9 | 3.8 | 1.4 | 173 | 168 | 5 | 3.9 | 22 |
| 127 | 0.4 | 0 | 0.009 | 1 | 308.4 | 31.2 | 3.2 | 1.6 | 195 | 187 | 8 | 4.7 | 29 |
| 128 | 0.4 | 0 | 0.009 | 1 | 347.8 | 30.3 | 3.2 | 1.6 | 207 | 202 | 5 | 8.9 | 54 |
| 129 | 0.4 | 0 | 0.009 | 1 | 240.6 | 21.5 | 1.2 | 1.4 | 150 | 145 | 5 | 3.9 | 19 |
| 130 | 0.4 | 0 | 0.009 | 1 | 235.6 | 16.6 | 4.8 | 1.4 | 143 | 137 | 6 | 3.7 | 16 |
| 131 | 0.7 | 0 | 0.009 | 1 | 565.7 | 7.9 | 2.9 | 1.6 | 454 | 449 | 5 | 15.8 | 66 |
| 132 | 0.7 | 0 | 0.009 | 1 | 730.6 | 12.1 | 2.1 | 1.4 | 556 | 551 | 5 | 18.8 | 91 |
| 133 | 0.7 | 0 | 0.009 | 1 | 642.1 | 14.3 | 3.5 | 1.4 | 517 | 510 | 7 | 15.2 | 73 |
| 134 | 0.7 | 0 | 0.009 | 1 | 398.8 | 7.9 | 1.5 | 1.6 | 326 | 322 | 4 | 15.3 | 63 |
| 135 | 0.7 | 0 | 0.009 | 1 | 771.3 | 8.8 | 8.2 | 1.7 | 582 | 572 | 10 | 24.5 | 129 |
| 136 | 0.7 | 0 | 0.009 | 1 | 356.8 | 6.9 | 6.6 | 1.5 | 291 | 282 | 9 | 8.8 | 44 |
| 137 | 0.7 | 0 | 0.009 | 1 | 424.9 | 16.3 | 2.2 | 1.5 | 326 | 320 | 6 | 12.5 | 77 |
| 138 | 0.7 | 0 | 0.009 | 1 | 369.3 | 16.2 | 1.3 | 1.4 | 281 | 274 | 7 | 7.4 | 33 |
| 139 | 0.7 | 0 | 0.009 | 1 | 367.2 | 18.8 | 3.1 | 1.4 | 287 | 281 | 6 | 14.6 | 89 |
| 140 | 0.7 | 0 | 0.009 | 1 | 352.2 | 10.4 | 4.4 | 1.3 | 274 | 269 | 5 | 10.1 | 45 |
| 141 | 0.4 | 0.3 | 0.009 | 1 | 138.3 | 14.8 | 1.2 | 1.6 | 78 | 74 | 4 | 1.9 | 10 |
| 142 | 0.4 | 0.3 | 0.009 | 1 | 105.2 | 16.1 | 5.8 | 1.3 | 75 | 66 | 9 | 2.5 | 12 |
| 143 | 0.4 | 0.3 | 0.009 | 1 | 136.9 | 28.0 | 2.4 | 1.4 | 77 | 72 | 5 | 3.7 | 17 |
| 144 | 0.4 | 0.3 | 0.009 | 1 | 110.6 | 26.9 | 4.8 | 1.3 | 71 | 62 | 9 | 1.5 | 8 |
| 145 | 0.4 | 0.3 | 0.009 | 1 | 107.8 | 22.4 | 1.8 | 1.7 | 62 | 57 | 5 | 1.5 | 7 |
| 146 | 0.4 | 0.3 | 0.009 | 1 | 75.5 | 28.7 | 4.5 | 1.3 | 46 | 41 | 5 | 1.4 | 9 |
| 147 | 0.4 | 0.3 | 0.009 | 1 | 105.5 | 27.0 | 4.1 | 1.4 | 63 | 56 | 7 | 1.4 | 7 |
| 148 | 0.4 | 0.3 | 0.009 | 1 | 47.7 | 18.5 | 6.2 | 1.1 | 42 | 31 | 11 | 0.7 | 5 |
| 149 | 0.4 | 0.3 | 0.009 | 1 | 70.9 | 30.6 | 1.0 | 1.4 | 40 | 37 | 3 | 1.1 | 5 |
| 150 | 0.4 | 0.3 | 0.009 | 1 | 64.7 | 25.9 | 5.6 | 1.2 | 48 | 39 | 9 | 1.1 | 7 |
| 151 | 0.7 | 0.3 | 0.009 | 1 | 120.9 | 4.7 | 4.0 | 1.4 | 108 | 101 | 7 | 4.1 | 19 |
| 152 | 0.7 | 0.3 | 0.009 | 1 | 283 | 6.7 | 3.0 | 1.4 | 210 | 204 | 6 | 7.5 | 44 |
| 153 | 0.7 | 0.3 | 0.009 | 1 | 201 | 9.7 | 2.2 | 1.6 | 165 | 160 | 5 | 6.2 | 25 |
| 154 | 0.7 | 0.3 | 0.009 | 1 | 164.2 | 8.8 | 6.8 | 1.2 | 141 | 133 | 8 | 4.9 | 19 |
| 155 | 0.7 | 0.3 | 0.009 | 1 | 132.0 | 14.8 | 0.7 | 1.4 | 97 | 96 | 1 | 3.5 | 20 |
| 156 | 0.7 | 0.3 | 0.009 | 1 | 321.9 | 9.4 | 1.4 | 1.7 | 224 | 220 | 4 | 10.2 | 43 |
| 157 | 0.7 | 0.3 | 0.009 | 1 | 314.6 | 16.7 | 1.6 | 1.5 | 203 | 200 | 3 | 8.1 | 35 |
| 158 | 0.7 | 0.3 | 0.009 | 1 | 135.5 | 11.2 | 1.9 | 1.4 | 101 | 95 | 6 | 4.2 | 23 |
| 159 | 0.7 | 0.3 | 0.009 | 1 | 164.9 | 11.2 | 2.1 | 1.3 | 127 | 124 | 3 | 5.4 | 22 |
| 160 | 0.7 | 0.3 | 0.009 | 1 | 194.7 | 13.7 | 2.8 | 1.5 | 146 | 140 | 6 | 4.1 | 17 |
